# Controlled Administration of Dehydrochloromethyltestosterone in Humans: Urinary Excretion and Long-Term Detection of Metabolites for Anti-Doping Purpose

**DOI:** 10.1101/2021.08.02.454719

**Authors:** Steffen Loke, Xavier de la Torre, Michele Iannone, Giuseppe La Piana, Nils Schlörer, Francesco Botrè, Matthias Bureik, Maria Kristina Parr

## Abstract

Dehydrochloromethyltestosterone (DHCMT) is an anabolic-androgenic steroid that was developed by Jenapharm in the 1960s and was marketed as Oral Turinabol^®^. It is prohibited in sports at all times; nevertheless, there are several findings by anti-doping laboratories every year. New long-term metabolites have been proposed in 2011/12, which resulted in adverse analytical findings in retests of the Olympic games of 2008 and 2012. However, no controlled administration trial monitoring these long-term metabolites was reported until now. In this study, DHCMT (5 mg, p.o.) was administered to five healthy male volunteers and their urine samples were collected for a total of 60 days. The unconjugated and the glucuronidated fraction were analyzed separately by gas chromatography coupled to tandem mass spectrometry. The formation of the described long-term metabolites was verified, and their excretion monitored in detail.

Due to interindividual differences there were several varieties in the excretion profiles among the volunteers. The metabolite M3, which has a fully reduced A-ring and modified D-ring structure, was identified by comparison with reference material as 4α-chloro-17β-hydroxymethyl-17α-methyl-18-nor-5α-androstan-13-en-3α-ol. It was found to be suitable as long-term marker for the intake of DHCMT in four of the volunteers. In one of the volunteers, it was detectable for 45 days after single oral dose administration. However, in two of the volunteers M5 (already published as long-term metabolite in the 1990s) showed longer detection windows. In one volunteer M3 was undetectable but another metabolite, M2, was found as the longest detectable metabolite.

The last sample clearly identified as positive was collected between 9.9 and 44.9 days. Furthermore, the metabolite epiM4 (partially reduced A-ring and a modified D-ring structure which is epimerized in position 17 compared to M3) was identified in the urine of all volunteers with the help of chemically synthesized reference as 4-chloro-17α-hydroxymethyl-17β-methyl-18-nor-androsta-4,13-dien-3β-ol. It may serve as additional confirmatory metabolite.

It is highly recommended to screen for all known metabolites in both fractions, glucuronidated and unconjugated, to improve identification of cheating athletes. This study also offers some deeper insights into the metabolism of DHCMT and of 17α-methyl steroids in general.

## Introduction

Dehydrochloromethyltestosterone (4-chloro-17β-hydroxy-17α-methyl-androsta-1,4-dien-3-one, DHCMT) is an anabolic-androgenic steroid which was introduced into the market as “Oral-Turinabol” by East German pharmaceutical company Jenapharm in the 1960s [1, 2]. It is a derivative of testosterone with enhanced anabolic properties which is orally bioavailable due to its alkylation at C-17 [3, 4]. According to Zafferoni *et al.* the chlorination at C-4 does not affect the anabolic or androgenic activity of DHCMT, which is contrary to the activity of progestogens or corticoids after substitution in this position [5]. However, the chloro-atom in position 4 hampers the reduction of the 4,5-double bond by 5α-reductase [6, 7]. 4-chlorination as well as the 1,2-double bond prevent the aromatization of the A-ring [4, 6]. Even if the androgenic activity is lower than that of testosterone, muscle tightness, acne, functional liver damage, and amenorrhea in women are reported as side effects [1, 8].

DHCMT was primarily developed to support recovery after major surgeries. However, shortly after its introduction on the market, it was misused in sports as performance-enhancing drug in one of the biggest systematic doping programs, until the German Democratic Republic’s collapse [1]. “Oral-Turinabol” was sold in two dosages, 1 and 5 mg, with recommendations for daily intake up to 50 mg [1]. It is explicitly mentioned as anabolic androgenic steroid (AAS) in class S1 in the list of prohibited substances published by the World Anti-Doping Agency (WADA) every year [9]. Even if discontinued as approved drug, the substance is still misused in sports until today. DHCMT gained new importance in doping control in the last 15 years with the number of its adverse analytical findings steadily increasing (Figure 1) through illegal marketing of steroid hormones mainly as so called “supplements” sold over the internet [10, 11]. Unfortunately, the control of these products is nearly impossible [12]. The WADA-accredited laboratories reported 108 adverse analytical findings for DHCMT in 2018 [13]. This fact is a clear indicator of the continuing relevance of research on substances even if they are on the market for a long time.

**Figure 1:**
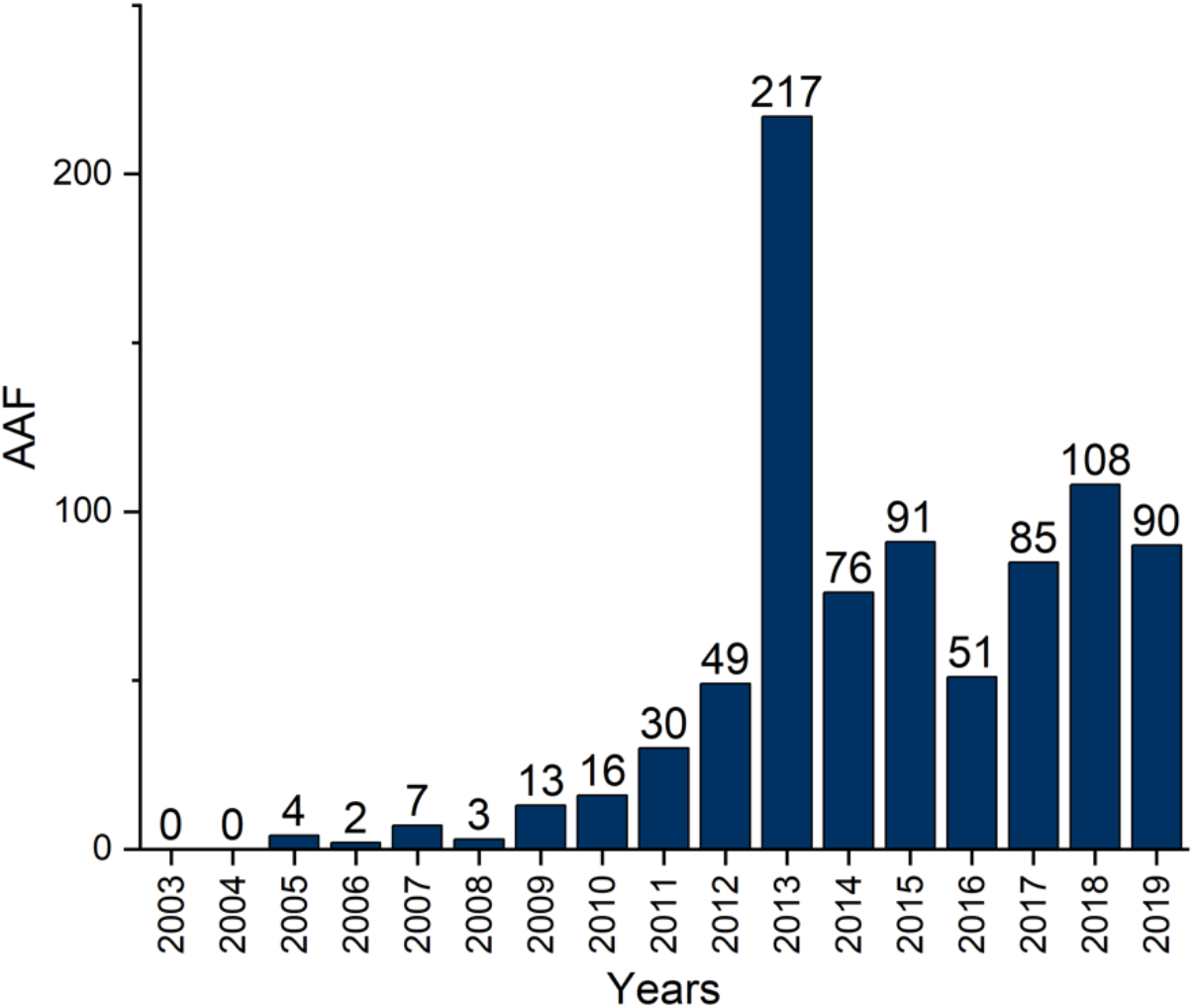
Adverse analytical findings of DHCMT between 2003 and 2019, according to [13]

Anabolic-androgenic steroids are extensively modified in the body by phase I and phase II metabolism. These reactions result in more polar substances facilitating the excretion via the urine [14, 15].

17β-Hydroxymethyl-17α-methyl-18-norandrost-13-ene metabolites of 17α-methyl steroids were first discovered in 2006, with the identification of a long-term metabolite of metandienone [16–18]. Subsequent research revealed analogous metabolites for other 17α-methyl steroids with capabilities for long-term detection, especially in case of oxandrolone and DHCMT [19–27].

As intermediates 17,17-dimethyl-18-norandrost-13-ene compounds are postulated. The subsequent hydroxylation at 17β-CH_3_ is catalyzed by steroid 21-hydroxylase (CYP21A2) [19, 27–30]. Continuing investigations on A-ring reduced long-term metabolites like 20ξOH-nortetrahydrochloromethyltestosterone (“M3”) by Sobolovsky and Rodchenkov led to several adverse analytical findings in retests of samples of the 2008 and 2012 Olympic Games [13, 31]. The underlying method of the research by Sobolovsky and Rodchenkov was based on the analysis of pooled urines from doping control samples already tested positive for DHCMT metabolites. As a controlled administration study was not yet reported, Kopylov *et al.* challenged these results of Sobolevsky [32]. In recent investigations the metabolite named “M3” in Sobolevsky’s publication was assigned to 4α-chloro-17β-hydroxymethyl-17α-methyl-18-nor-5α-androst-13-en-3α-ol by comparison with synthesized reference material by Kratena *et al.* [33]. Sobolevsky’s “M4” was analogously assigned to 4-chloro-17α-hydroxymethyl-17β-methyl-18-norandrosta-4,13-dien-3β-ol by our group in 2019 [26] and confirmed by Kratena *et al.* [25].

The pathways of DHCMT metabolism are not fully uncovered yet, but due to the work of Liu *et al.* and Stoll *et al.* [27, 29] some enzymes and metabolic processes came to the fore.

The trial reported in this manuscript aimed to monitor metabolite excretion after the intake of DHCMT based on a controlled administration trial in five male volunteers. Information about the long-term metabolism of DHCMT may help to further improve the fight against doping, support anti-doping laboratories with information on best target analytes and their detection windows. Furthermore, knowledge on biotransformation of 17β-hydroxy-17α-methyl-androstan-3-one steroids in general is increased.

## Materials and Methods

### Chemicals and Reagents

Methanol (HPLC-grade), t-butyl methyl ether (TBME, HPLC-grade), ethyl acetate, K_2_CO_3_, and NaH_2_PO_4_ were purchased from Merck KGaA (Darmstadt, Germany). KHCO3 was provided by Honeywell Fluka (Charlotte, USA). Na_2_HPO_4_ was bought from Carlo Erba (Val de Reuil, France). β-Glucuronidase was purchased from Roche Diagnostics (Mannheim, Germany), β-Glucuronidase from Helix Pomatia (G0876) and sulfatase from Helix Pomatia (S9626) were provided by Sigma Aldrich (Milano, Italy). Hydrochloric acid and acetic acid were provided by Fisher Scientific (Loughborough, United Kingdom). 17α-methyltestosterone was delivered by Steraloids (Newport, R.I., USA). Etiocholanolone-D_5_, androsterone-D_4_-glucuronide, and 4-chloro-6β,17β-dihydroxy-17α-methyl-androst-1,4-dien-3-one are coming from the National Measurement Institute (West Lindfield, New South Wales, Australia). N-Methyl-N-(trimethylsilyl)trifluoroacetamide (MSTFA) was bought from Chemische Fabrik Karl Bucher GmbH (Waldstetten, Germany). Dehydrochloromethyltestosterone (DHCMT) was obtained from Cohnchem Scientific Co., Ltd (Derby, UK). 4α-Chloro-17β-hydroxymethyl-17α-methyl-18-nor-5α-androst-13-en-3α-ol was obtained from the Austrian Anti-Doping Laboratory, provided by the World Association of Anti-Doping Scientists (WAADS). All other chemicals were purchased from Sigma-Aldrich (Munich, Germany).

### GC-QQQ-MS analysis

The gas chromatographic-mass spectrometric analysis of urine samples was performed on an Agilent 7890A gas chromatographic system coupled to an Agilent 7000 GC/MS triple quadrupol (QQQ) mass spectrometer with the following parameters for the analysis of the intermediates and products as reported by Martinez-Brito et al. [34]: Agilent HP1 (17 m, 0.20 mm, 0.11 μm), carrier gas: helium, oven program: 188 °C, hold for 2.5 min, +3 °C/min to 211 °C, hold for 2.0 min, +10 °C/min to 238 °C, +40 °C to 320 °C, hold for 3.2 min, injection volume: 2 μL, split: 20:1, injection temperature: 280 °C, electron ionization (EI): 70 eV, transitions, collision energy and retention time for each substance are displayed in Table 1.

**Table 1:** Overview of screened compounds including the retention time, the transitions and the collision energy used for analysis

| Substance | Retention time [min] | Transition | CE [eV] |
| --- | --- | --- | --- |
| DHCMT | 16.40 | 478.4 → 240 | 5 |
|  |  | 478.4 → 225 | 5 |
|  |  | 240 → 225 | 5 |
|  |  | 240 → 189 | 5 |
|  |  | 225 → 189 | 5 |
| M1 | 16.68 | 315 → 227 | 10 |
|  |  | 227 → 163 | 5 |
|  |  | 227 → 93 | 5 |
| M2 | 14.50 | 377 → 251 | 15 |
|  |  | 377 → 185 | 22 |
|  |  | 287 → 251 | 15 |
|  |  | 287 → 185 | 12 |
| M3 | 15.25 | 381 → 343 | 15 |
|  |  | 381 → 253 | 15 |
|  |  | 379 → 343 | 7 |
|  |  | 379 → 253 | 13 |
|  |  | 379 → 147 | 15 |
| epiM3 | 14.40 | 381 → 343 | 15 |
|  |  | 381 → 253 | 15 |
|  |  | 379 → 343 | 7 |
|  |  | 379 → 253 | 10 |
|  |  | 379 → 147 | 15 |
| M4 | 14.30 | 379 → 149 | 15 |
|  |  | 377 → 149 | 13 |
|  |  | 287 → 125 | 22 |
| epiM4 | 15.10 | 379 → 149 | 15 |
|  |  | 377 → 149 | 13 |
|  |  | 287 → 125 | 22 |
| M5 | 16.60 | 658.6 → 244 | 15 |
|  |  | 658.6 → 244 | 17 |

Every sample was treated with 50 μL of TMIS reagent (MSTFA / ammonium iodide / ethanethiol, 1000:2:3, v:w:v) at 75 °C for 20 min prior to injection to generate the per-TMS derivatives.

### GC-QTOF-MS analysis

High resolution accurate mass analyses were performed on an Agilent GC-quadrupol time-of-flight (QToF) 7890B/7250 (Agilent Technologies, Milano, Italy), equipped with an Agilent HP1 column (17 m, 0.20 mm, 0.11 μm) with helium as carrier gas as reported earlier [35, 36]. Injection was performed in split mode with a 1:10 ratio at 280 °C. The oven program had the following heating rates: 188 °C hold for 2.5 min, 3 °C/min to 211 °C and hold for 2 min, 10 °C/min to 238 °C, 40 °C/min to 320 °C and hold for 3.2 min. The hyphenated QToF was operated in full scan with an ionization energy of 70 eV. Aberrantly, in LEI an ionization energy of 15 eV. Ions were detected from *m/z* 50 to 750.

### GC-MS analysis

The gas chromatographic-mass spectrometric analysis of the extended steroid profile samples was performed on an Agilent 6890N gas chromatographic system coupled to an Agilent 5973N GC/MSD with the following parameters for the analysis of the intermediates and products: Agilent HP5 (17 m, 0.20 mm, 0.11 μm), carrier gas: helium, oven program: 60 °C, hold for 0.5 min, +60 °C/min to 210 °C, +2.5 °C/min to 270 °C, +30 °C to 320 °C, hold for 2.5 min, injection volume: 2 μL, split: 20:1, injection temperature: 280 °C, electron ionization (EI): 70 eV

### Nuclear Magnetic Resonance

The nuclear magnetic resonance (NMR) analyses were performed at 500 MHz (^1^H NMR) and 125 MHz (^13^C NMR) at 296 K on a Bruker (Rheinstetten, Germany) Avance III 500 instrument equipped with a nitrogen-cooled 5 mm inverse probe head (Prodigy TCI) with actively shielded z-gradient coil. Chemical shifts are reported in δ values (ppm) relative to tetramethylsilane. Solutions of about 5 mg of each compound in deuterated dimethylsulfoxid (d_6_-DMSO) were used for conducting ^1^H; H,H COSY; ^13^C{^1^H} APT; H,C HMQC; H,C HMBC and H,H NOESY experiments. 2D experiments were acquired applying non-uniform sampling (NUS, [37]).

### Synthesis of Reference Substances

4-Chloro-17α-hydroxymethyl-17β-methyl-18-norandrosta-4,13-dien-3β-ol and three of its diastereomers were synthesized as described in the supplemental information. After purification analytical characterization was performed using GC-QTOF-MS and NMR analyses.

### Preparative Liquid Chromatography

The preliminary purification runs were performed on ISOLERA ONE (Biotage AB, Uppsala, Sweden) instrument equipped with UV-Vis detector (detection wavelength 254 nm with 15 mAU as threshold). The column used was a BIOTAGE SNAP (Biotage AB, Uppsala, Sweden) Ultra (10 g, 25 μm). The flow used was 12 mL/min with an isocratic elution with solvent composition of hexane:EtOAc (60:40, v:v).

### HPLC Separation

The HPLC runs were performed on Agilent 1260 instrument equipped with C18 (250 mm x 10 mm, particle size 5 μm) THERMO electron corporation (Waltham, Massachusetts, USA) column. The flow was 2.5 mL, the maximum flow gradient was 1 mL/min. An isocratic run with solvent composition MeOH:H2O (70:30, v:v) was performed. The injection volume was 0.5 mL. The selected wavelength for the UV detection were 250 nm, 254 nm and 210 nm.

### Human Administration Trial

An administration study with a single oral intake of 5 mg DHCMT was performed in five healthy male volunteers (age: 30 - 67 years). The study was approved by the ethics committee of the School of Pharmaceutical Science and Technology, Tianjin University. The study was performed following the recommendations of the Helsinki declaration and written informed consent was obtained from the participants. The anthropometric data and further characteristics of the volunteers are shown in Table 2.

**Table 2:**
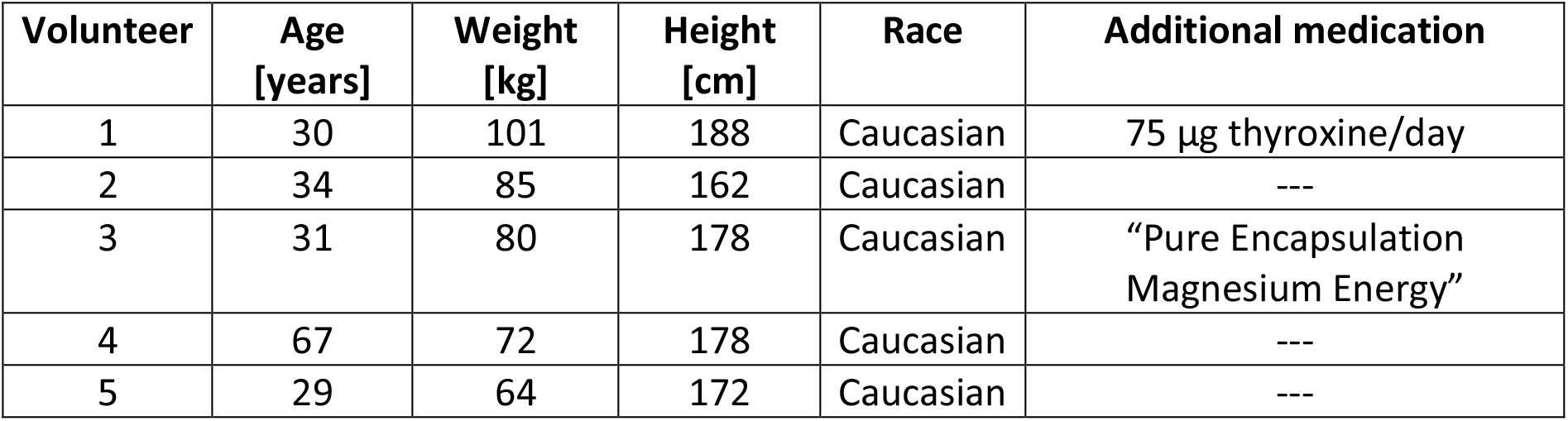
Characteristics of the volunteers

| Volunteer | Age [years] | Weight [kg] | Height [cm] | Race | Additional medication |
| --- | --- | --- | --- | --- | --- |
| 1 | 30 | 101 | 188 | Caucasian | 75 µg thyroxine/day |
| 2 | 34 | 85 | 162 | Caucasian | --- |
| 3 | 31 | 80 | 178 | Caucasian | “Pure Encapsulation Magnesium Energy” |
| 4 | 67 | 72 | 178 | Caucasian | --- |
| 5 | 29 | 64 | 172 | Caucasian | --- |

Blank urines were collected within 24 hours before the administration of DHCMT. During the first 72 hours after intake, every urine was collected. Afterwards, every first urine in the morning was collected until day 60 post administration. The urines were directly frozen after every collection and kept at −24 °C until analysis. Specific gravity for each sample was determined with two drops of urine on a Mettler Toledo RE50 refractometer prior to analysis. The correction factor was calculated by (1,020-1,000)/(SG-1,000). SG is the specific gravity of the sample and 1,020 is the value to which the specific gravity of urine is normalized.

### Sample Preparation for Metabolite Analysis

Samples were analyzed using the method adapted from the Anti-Doping Laboratory Rome for DHCMT metabolite confirmation. Briefly, an aliquot of 6 mL was used for the following analysis. After the addition of an internal standard (50 μL solution of methyltestosterone [100 μg/mL]), the urine was extracted with 10 mL of TBME. The organic layer was separated and evaporated to dryness to result in the fraction of unconjugated metabolites (free fraction).

For the extraction of the glucuronide fraction, the aqueous layer was used. The internal standard (50 μL solution of methyltestosterone [100 μg/mL]) and a standard containing deuterated glucuronides (50 μL solution of a mixture of androsterone-D_4_-glucuronide, etiocholanolone-D_5_; 24 μg/mL) were added. After the addition of 750 μL of phosphate buffer (0.8 M; 71.2 g Na_2_HPO_4_, 55.0 g NaH_2_PO_4_, ad 1000 mL H_2_O) and 50 μL β-glucuronidase, the mixture was incubated at 55 °C for 60 min. Afterward, 500 μL of carbonate/bicarbonate buffer (20 %; 200 g K_2_CO_3_, 200 g KHCO_3_, ad 1000 mL H_2_O) was added and the mixture was extracted with 10 mL of TBME. The ether layer was evaporated to dryness. Analyses were performed by GC-QQQ-MS and GC-QTOF-MS.

A standard containing 90 μL of a solution of DHCMT, metabolite M1, and metabolite M3 (1 μg/mL each), and 9 μL epiM4 (100 μg/mL), and 50 μL methyltestosterone (100 μg/mL) was evaporated to dryness. This mixture was measured together with every batch of samples. The chromatogram of this mixture is displayed in Figure 2.

**Figure 2:**
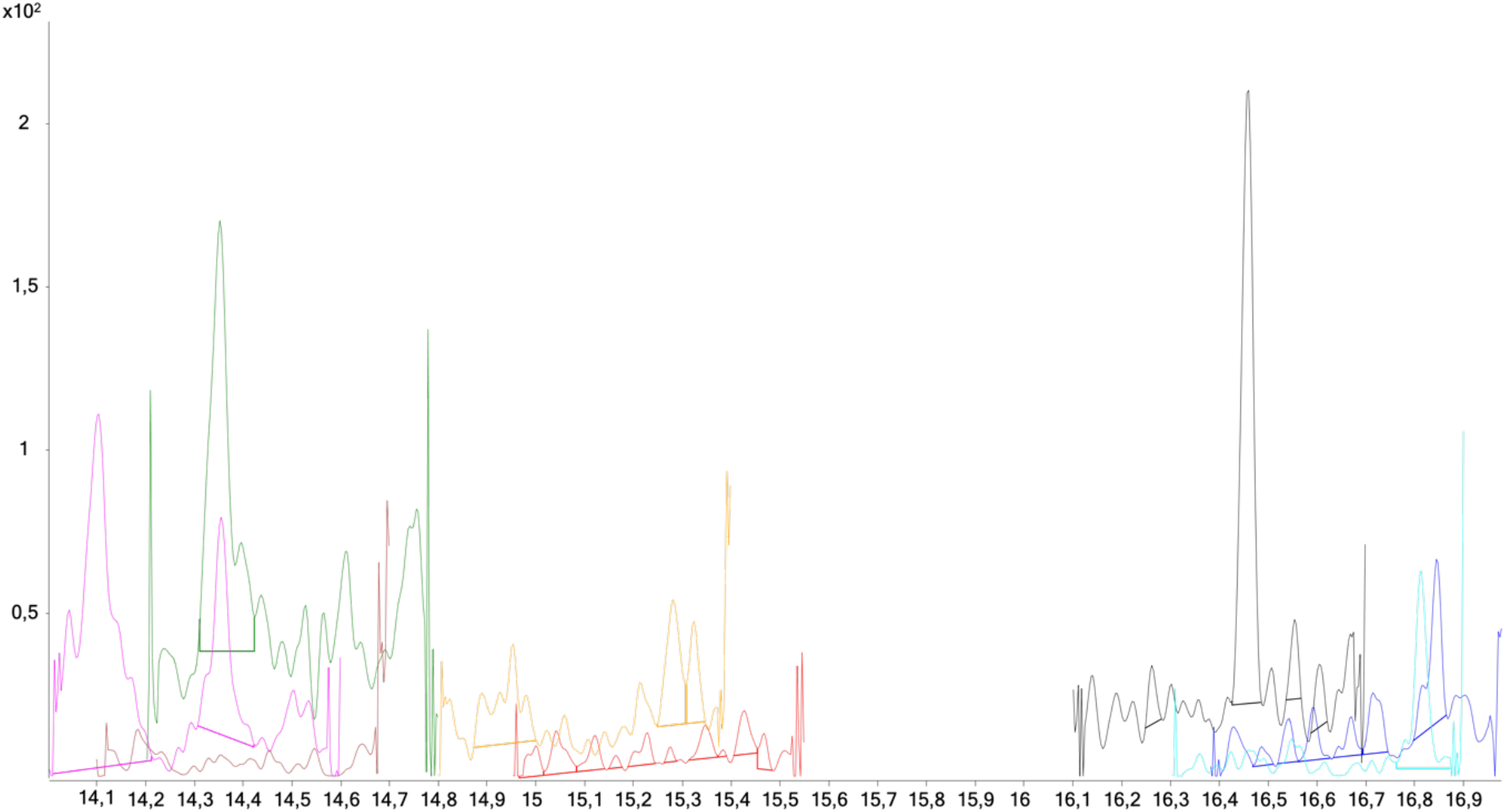
GC-QQQ-MS chromatogram of the standard mixture; yellow: epiM4, red: M3, black: parent compound, blue: M1

### Extended Steroid Profile

Solid-phase extraction was performed using Waters Sep-Pak Classic C_18_ cartridges (WAT051910, Waters Corporation, Massachusetts, USA). The cartridge was conditioned with 2 x 2 mL of each methanol and water, loaded with 2 mL of blank urine, washed with 2 x 2 mL of water, and eluted with 2 x 2 mL of methanol. The combined eluates were evaporated to dryness. After adding β-glucuronidase and sulfatase from *Helix pomatia* in acetate buffer, the mixture was incubated at 55 °C for 3 h. Afterwards, 500 μL of carbonate buffer was added, and the mixture was extracted with 5 mL TBME, and an internal standard was added (50 μL of a mixture of 5α-androstane-3α,17α-diol, stigmasterol, cholesterol butyrate, androsterone-D_4_, etiocholanolone-D_5_-sulfate, cortisol-D4; concentration: 5 μg/mL each). The ether layer was evaporated to dryness. The derivatization was performed in three steps: after the addition of 100 μL of N-methylhydroxylamine hydrochloride in pyridine (2 %), the sample was heated to 55 °C for 1 hour, the solvent was evaporated afterwards. Following the addition of 50 μL 1-(trimethylsilyl)imidazole, the sample was heated with a microwave oven, extracted with cyclohexane, and evaporated to dryness; finally, 50 μL MSTFA were added, and the sample was heated to 75 °C for 10 minutes. Analysis of the derivatized residues was performed by GC-MS.

## Results

### Metabolite Detection

The analysis mainly focused on metabolites Sobolevsky *et al.* proposed in 2012 [23] as target analytes. Analysis was performed separately for the unconjugated and for the glucuronidated metabolites. Structure assignments of the aglycons extracted in the different fractions (with or without deglucurinidation) are based on the comparison with authentic reference material in case of M1, M3 and epiM4 as described below.

The success of the deglucuronidation was evaluated by comparison of two substances in the added standard: etiocholanolone-[D_5_] and androsterone-[D_4_]-glucuronide. The mean value of deglucuronidation in all volunteers was 59.0 %.

Most of the compounds were only excreted as glucuronides (M2, M3, M4, epiM4) and thus detectable in the extract after deconjugation (glucuronide fraction). The other substances (DHCMT, M1, M5) were found in both fractions (unconjugated and glucuronide fraction). The metabolite epiM3 could not be confirmed in any sample but urines of volunteer 5 gave some samples with signals different to the blank urine between 1.7 and 7.0 days. Figure 3 gives an overview of the found metabolites in comparison to a blank sample.

**Figure 3:**
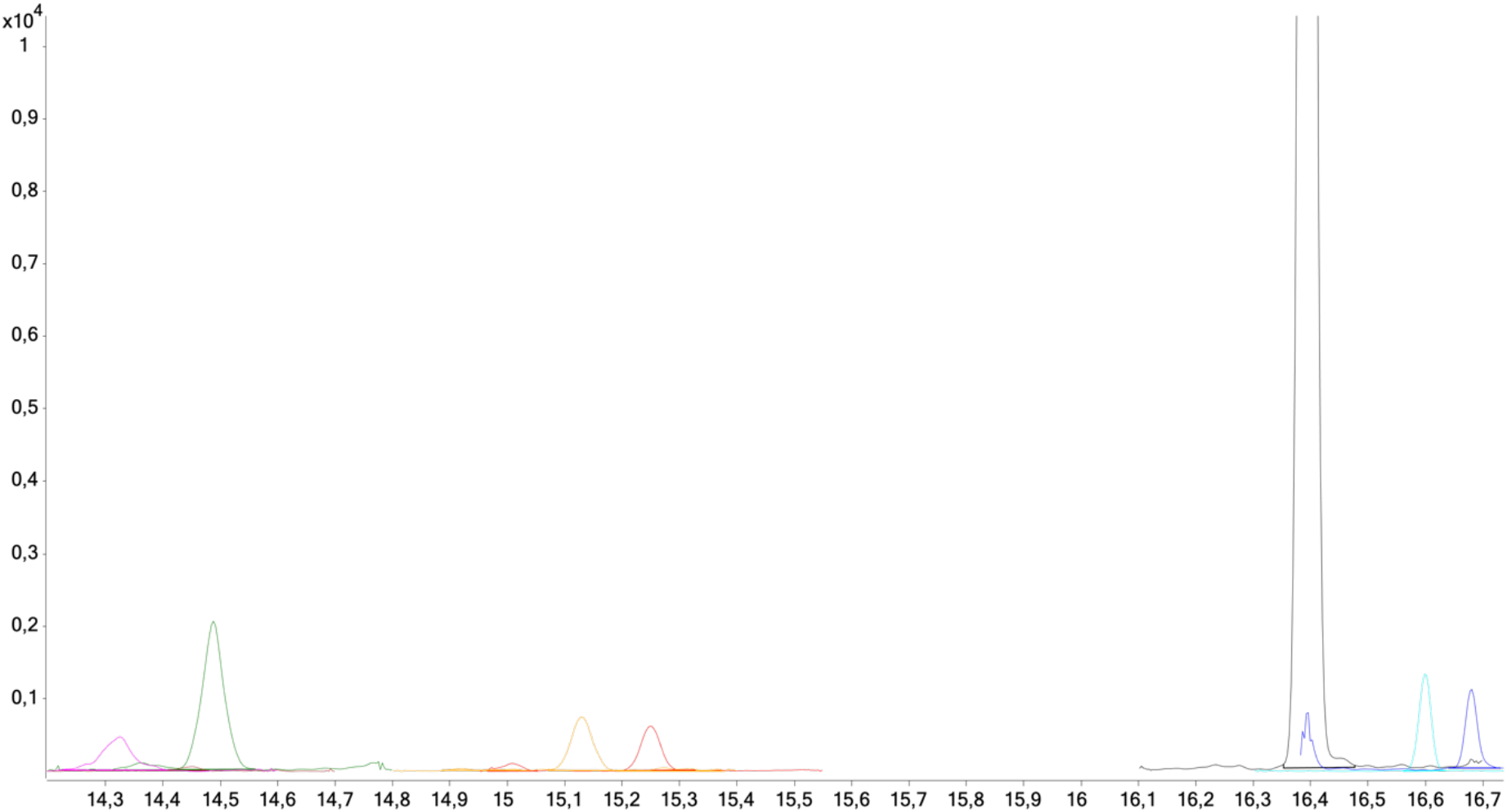
upper picture: GC-QQQ-MS chromatogram of a blank sample of volunteer 1; lower picture: GC-QQQ-MS chromatogram of a post-administration sample of volunteer 1 (25.3 h after intake); magenta: M4, green: M2, yellow: epiM4, red: M3, black: parent compound, turquoise: M5, blue: M1

Aberrantly from the other volunteers, volunteer 4 displayed a completely different excretion profile. Neither M3 nor M4 were detected in his samples. The last sample with a positive metabolite finding was collected after 18 days. The metabolite detected was M2.

### Synthesis of 4-Chloro-17α-hydroxymethyl-17β-methyl-18-nor-androsta-4,13-dien-3β-ol

For proper structure confirmation of M4/epiM4, different diastereomers of 4-chloro-17α-hydroxymethyl-17β-methyl-18-nor-androsta-4,13-dien-3β-ol were synthesized starting from 4-chloro-androst-4-ene-3,17-dione. The synthesis route is described in the supplement. After the regioselective reduction of the 3-oxo group, the protection of the 3-hydroxy group followed and the intermediate was transformed using a method adapted from Kratena *et al.* [25]. A new carbon-atom at position 17 was introduced. After epoxidation of the 17(20)-double bond, the epoxide was opened under acidic conditions which went along with a Wagner-Meerwein rearrangement mainly resulting in 17α-hydroxymethyl-17β-methyl derivatives. This is called epiM4 following Sobolevsky’s nomenclature that 17α-hydroxymethyl-17β-methyl derivatives are called “epi” whereas the 17β-hydroxymethyl-17α-methyl metabolites are described as, e.g., “M4”. The final step was the acid catalyzed deprotection of the 3-hydroxy group. Four diastereomeric products were obtained. The detailed steps are described in the supplement. The retention times and relative abundances are displayed in Table 3. As expected, their mass spectra are very similar. As an example, the spectrum of the main product, 4-chloro-17α-hydroxymethyl-17β-methyl-18-nor-androstane-4,13-dien-3β-ol, is displayed in Figure 4.

**Table 3:**
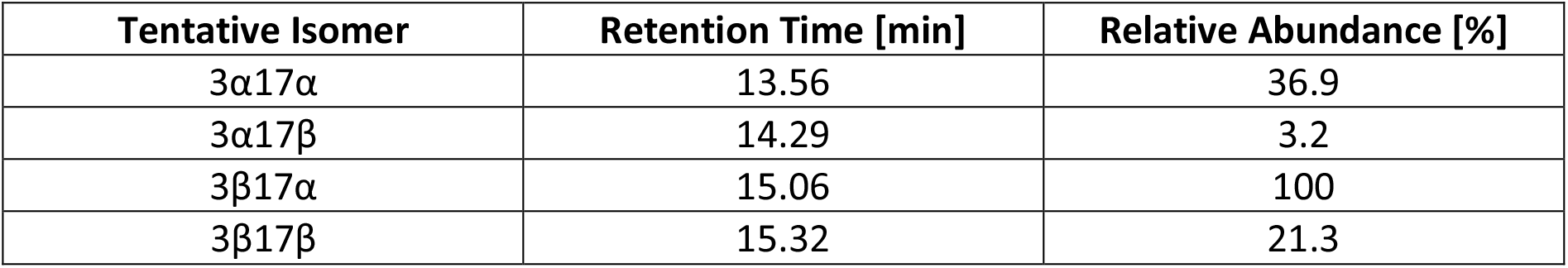
Retention times and relative abundances of the 4-chloro-17ξ-hydroxymethyl-17ξ-methy-18-norl-androstane-4,13-dien-3ξ-ol isomers

**Figure 4:**
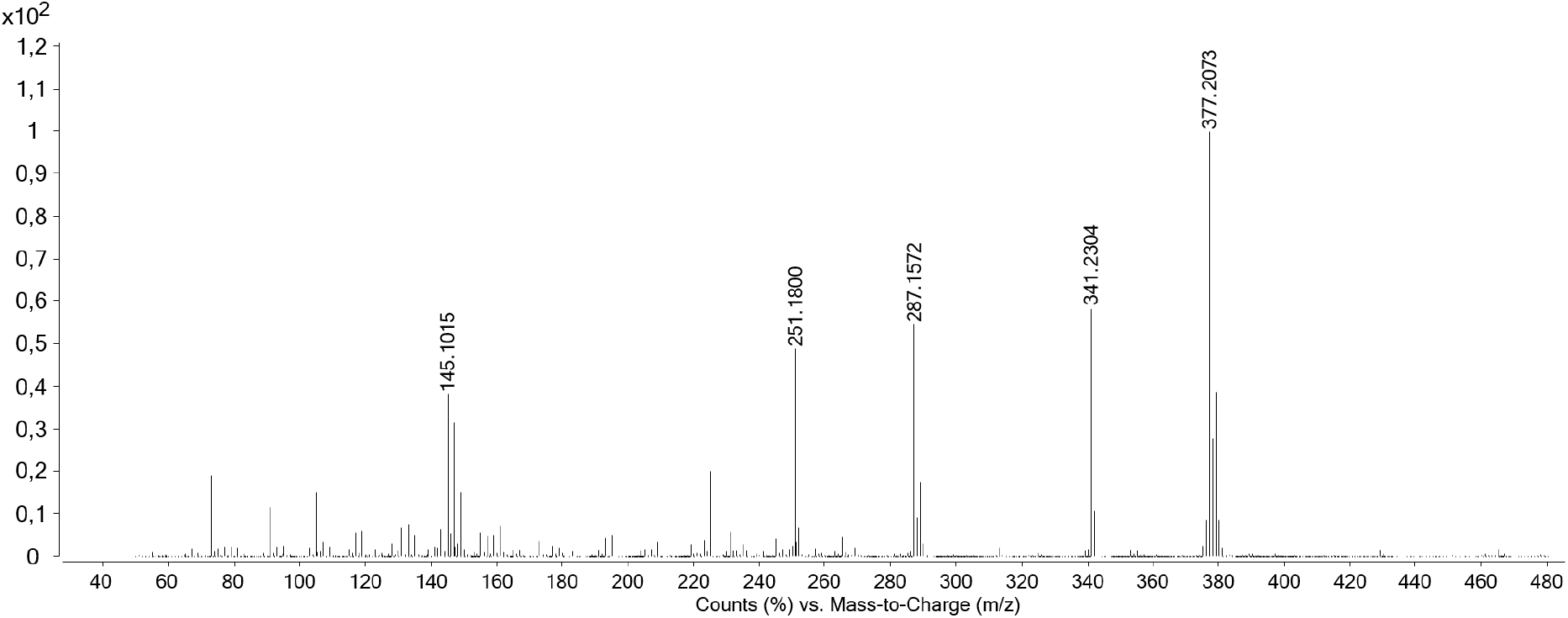
Mass spectrum (GC-EI-QTOF-MS, 70 eV) of 4-chloro-17α-hydroxymethyl-17β-methyl-18-nor-androstane-4,13-dien-3β-ol (bis-trimethylsilyl)

The obtained main products had the 4-chloro-17α-hydroxymethyl-17β-methyl-18-nor-androstane-4,13-dien-3ξ-ol structure with the 3β-ol as major product. The 4-chloro-17β-hydroxymethyl-17α-methyl-18-nor-androstane-4,13-dien-3ξ-ol structures appeared as minor byproducts.

After purification of the major diastereomer by liquid chromatography, structure confirmation by NMR was achieved. All signals were assigned from shifts and correlations of 1D and 2D measurements:

^1^H NMR (500 MHz, DMSO-d_6_): δ = 5.11 (d, ^3^J = 6.7 Hz, 1H, C-3-<u>OH</u>); 4.31 (t, ^3^J = 5.5 Hz, 1H, C-17-CH_2_-<u>OH</u>); 3.92 (ddm, ^3^J = 13.3/6.7 Hz, 1H, H-3a); 3.14 (d, ^3^J = 5.5 Hz, 2H, C-17-<u>CH_2_</u>-OH); 2.88 (ddd, ^3^J = 13.8/3.7/2.5 Hz, 1H, H-6a); 2.19 (m, 1H, H-15a); 2.14 (m, 1H, H-8); 2.04 (m, 1H, H-15b); 2.00 (m, 1H, H-6b); 1.95 (m, 1H, H-12b); 1.94 (m, 1H, H-7b); 1.91 (m, 1H, H-2a); 1.86 (m, 1H, H-16a); 1.80 (m, 1H, H-12a); 1.78 (m, 1H, H-11a); 1.74 (m, 1H, H-1b); 1.56 (m, 1H, H-2b); 1.36 (m, 1H, H-16b); 1.32 (m, 1H, H-1a); 1.12 (m, 1H, H-11b); 1.01 (s, 3H, C-10-<u>CH_3_</u>); 1.00 (m, 1H, H-9); 0.90 (s, 3H, C-17-<u>CH_3_</u>); 0.84 (m, 1H, H-7a).

^13^C NMR (125 MHz, DMSO-d_6_): δ = 141.4 (s, C-5), 138.8 (s, C-13), 137.4 (s, C-14), 130.4 (s, C-4), 68.7 (d, C-3), 67.9 (t, C-17-<u>CH_2_</u>-OH), 51.8 (d, C-9), 51.4 (s, C-17), 40.3 (s, C-10), 36.6 (d, C-8), 34.2 (t, C-16), 33.5 (t, C-1), 31.2 (t, C-7), 29.9 (q, C-15), 29.6 (t, C-2), 27.4 (t, C-6), 23.0 (t, C-12), 22.9 (t, C-11), 22.1 (q, C-17-<u>CH_3_</u>), 19.1 (q, C-10-<u>CH_3_</u>).

### Metabolite Structure Assignment

The targeted metabolites are different in their structures. Tentative structures have been proposed by Sobolevsky *et al.* [23] and Schänzer *et al.* [38].

The first metabolite, M1, is a monohydroxylated derivative of the parent compound. Its structure was confirmed as 6β-hydroxy-DHCMT by comparison of retention time and ion transitions with commercial reference material.

The metabolite M5 is also hydroxylated in position 6 but also functionalized in position 16 (16-oxo) and reduced in positions 3 (3-hydroxy) and 4(5) (no double bond). Its structure was proposed as 3α,6β,17β-trihydroxy-17α-methyl-4ξ-chloro-5β-androst-1-en-16-one by Schänzer *et al.* [38] based on MS fragmentations. To the best of our knowledge, no reference material for confirmation is available until now.

The other screened metabolites (M2, M3/epiM3 and M4/epiM4) have a rearranged D-ring which goes together with a shift of the 18-methyl group, concomitant with formation of a 13(14) double bond and hydroxylation at 17-CH_3_. As proposed by Sobolevsky *et al.* [23] the metabolites M2, M3/epiM3 and M4/epiM4 share the reduction in position 3 (3-hydroxy). The metabolites M3/epiM3 have a fully reduced A-ring. The structure of M3 was further elucidated by comparison with the authentic reference material of 4α-chloro-17β-hydroxymethyl-17α-methyl-18-nor-5α-androst-13-en-3α-ol, with matching retention time and ion transitions according to WADA criteria [39]. Supported by elution order epiM3 is supposed to be the 17-epimer of M3 [19, 28]. Metabolites M2 and M4/epiM4 still contain one double bond in the A-ring. While metabolite M2 is assigned to the 1-ene isomer, metabolites M4/epiM4 are supposedly reduced in position 1(2). Comparison with the inhouse synthesized reference confirmed epiM4 as 4-chloro-17α-hydroxymethyl-17β-methyl-18-nor-androstane-4,13-dien-3β-ol, while M4 is assigned to 4-chloro-17β-hydroxymethyl-17α-methyl-18-nor-androstane-4,13-dien-3α-ol with matching retention times and ion transitions in GC-QQQ-MS.

### Excretion Profiles of Metabolites

The excretion period is profoundly different depending on the fraction, the volunteer, or the compound itself. The given signal area was adjusted by the area of the internal standard and the measured specific gravity. Figure 5 displays the excretion kinetics of DHCMT from the glucuronide fraction.

**Figure 5:**
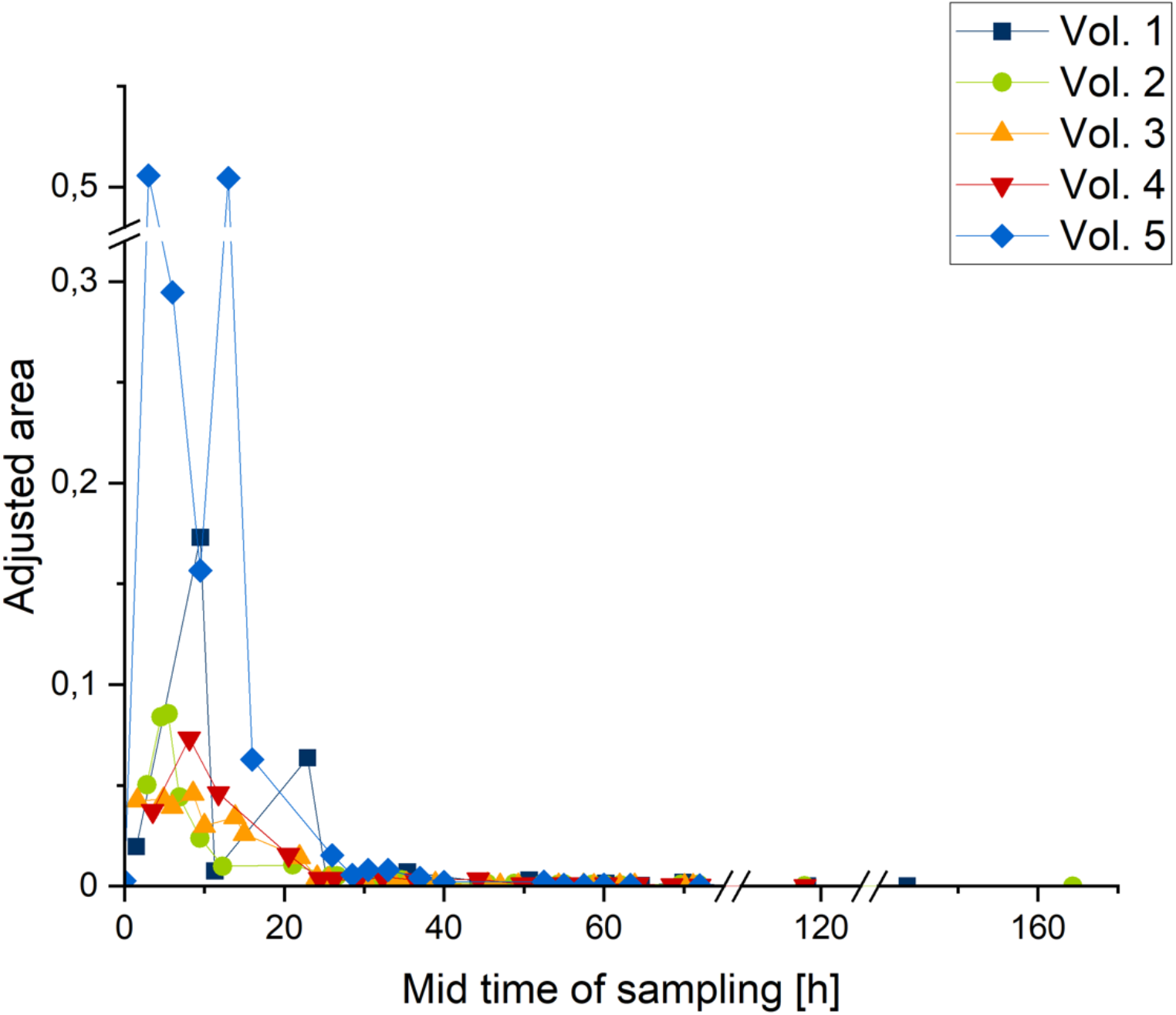
Elimination profile of parent compound DHCMT (detected in glucuronide fraction); peak area adjusted by internal standard and specific gravity

The parent compound (DHCMT) was detected in the unconjugated fraction up to 72 hours after the oral intake. In the glucuronide fraction it was found for a longer time (up to 6.9 days, volunteer 2). The maximum concentration was detected in the first few hours (midtime of sampling up to 10 hours post administration). Volunteers 1 and 5 had a distinct second peak in the excretion of the unconjugated DHCMT as well as of the conjugated substance (Figure 5). The above-mentioned fact of two peaks in the excretion profile is visible throughout all metabolites and volunteers, but it is remarkable only in volunteers 1 and 5.

The total amounts over all positive samples are displayed in Figure 6. The relative amount of DHCMT varies from 1.013 % (volunteer 4) and 7.127 % (volunteer 1) for the glucuronidated and between 0.009 % (volunteer 3) and 0.037 % (volunteer 1) for the unconjugated substance of the administered dose of 5 mg.

**Figure 6:**
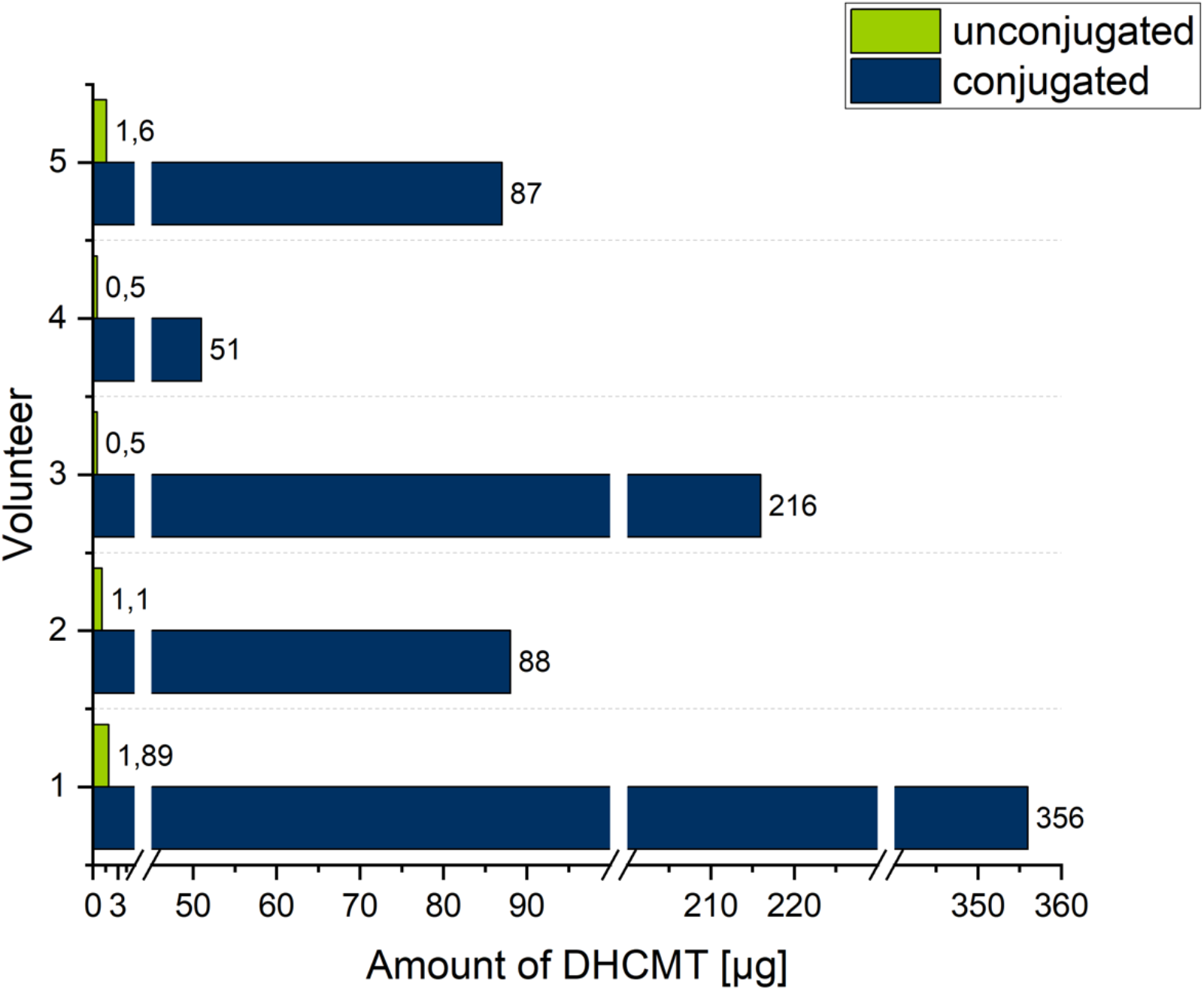
Total amount of excreted DHCMT as conjugated and unconjugated substance in μg

As mentioned above, three of the targeted compounds are found in both fractions. Besides the parent compound, the detection window of two metabolites, M1 and M5, was significantly longer in the unconjugated fraction than in the glucuronide fraction: M1 between 11 h (volunteer 5; glucuronide fraction: 26 h, free fraction: 37 h) and 48.2 h (volunteer 4; glucuronide fraction: 44,3 h, free fraction: 92,5 h), M5 between 44.0 h (volunteer 5; glucuronide fraction: 194 h, free fraction: 240 h) and 144.1 h (volunteer 1; glucuronide fraction: 213.1 h, free fraction: 357.2 h) longer. The detection times of all volunteers for both metabolites in the free and glucuronide fraction are displayed in Figure 7.

**Figure 7:**
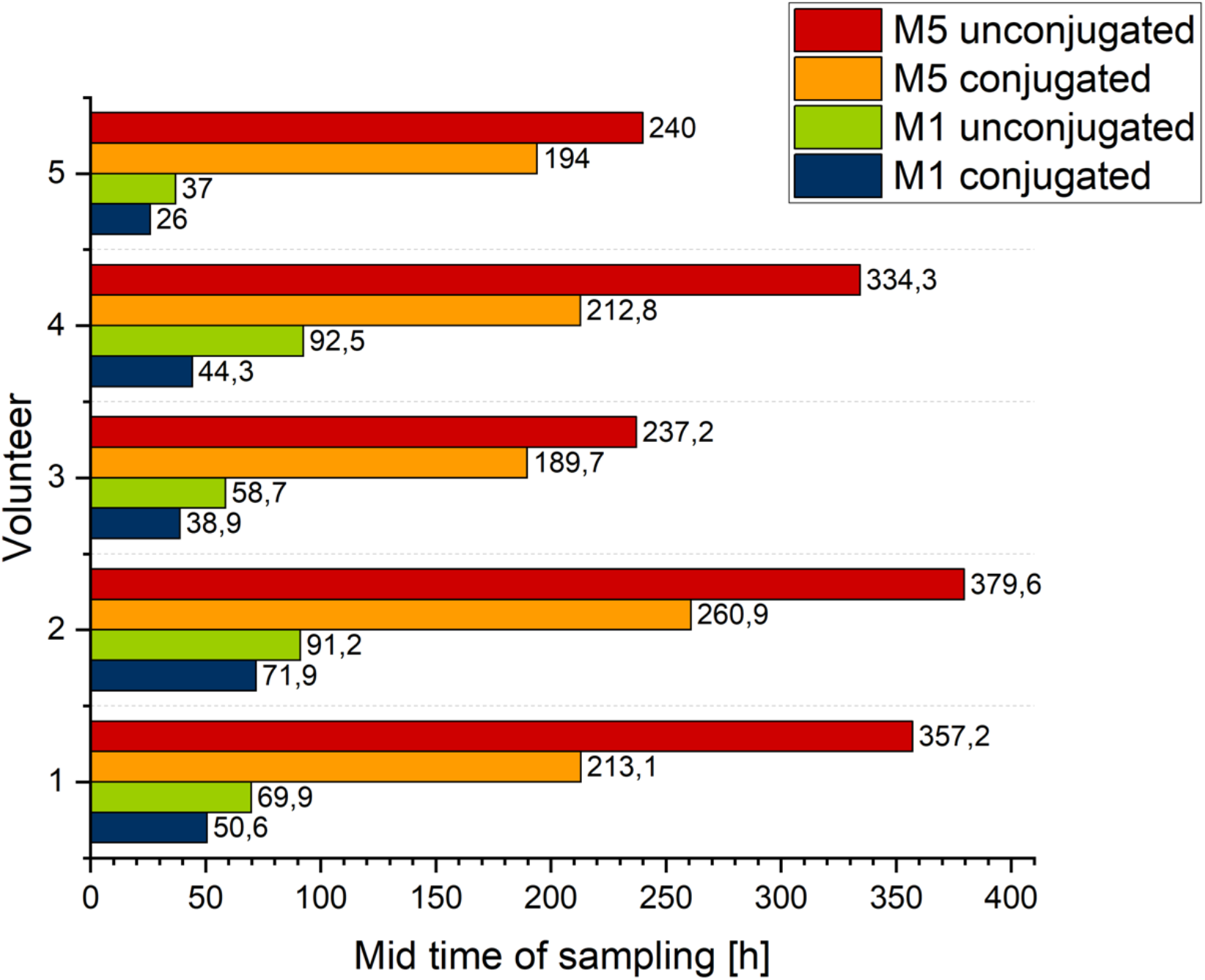
Comparison of the excretion times of metabolites M1 and M5 in the free vs. the glucuronide fraction (all volunteers)

The longest detectable metabolite of every volunteer (separately for unconjugated and glucuronide fraction) for each volunteer is displayed in Figure 8. The metabolite, which is excreted for the longest time, is different between the volunteers. Volunteer 1 excreted M3 for 44.9 days and volunteer 5 for 19.0 days in the glucuronide fraction. In contrast, volunteers 2 and 3 excreted the unconjugated M5 over 15.8 days and 9.9 days whereas M3 was only detectable for 7.8 and 4.9 days (Figure 9). It is noticeable that the later samples of volunteers 1, 2, 3, and 5 had signals different to the blank urine for the metabolite M3 (vol. 1: 58.9 days; vol. 2: 58.8 days; vol. 3: 56.9 days; vol. 5: 57.1 days; Figure 10).

**Figure 8:**
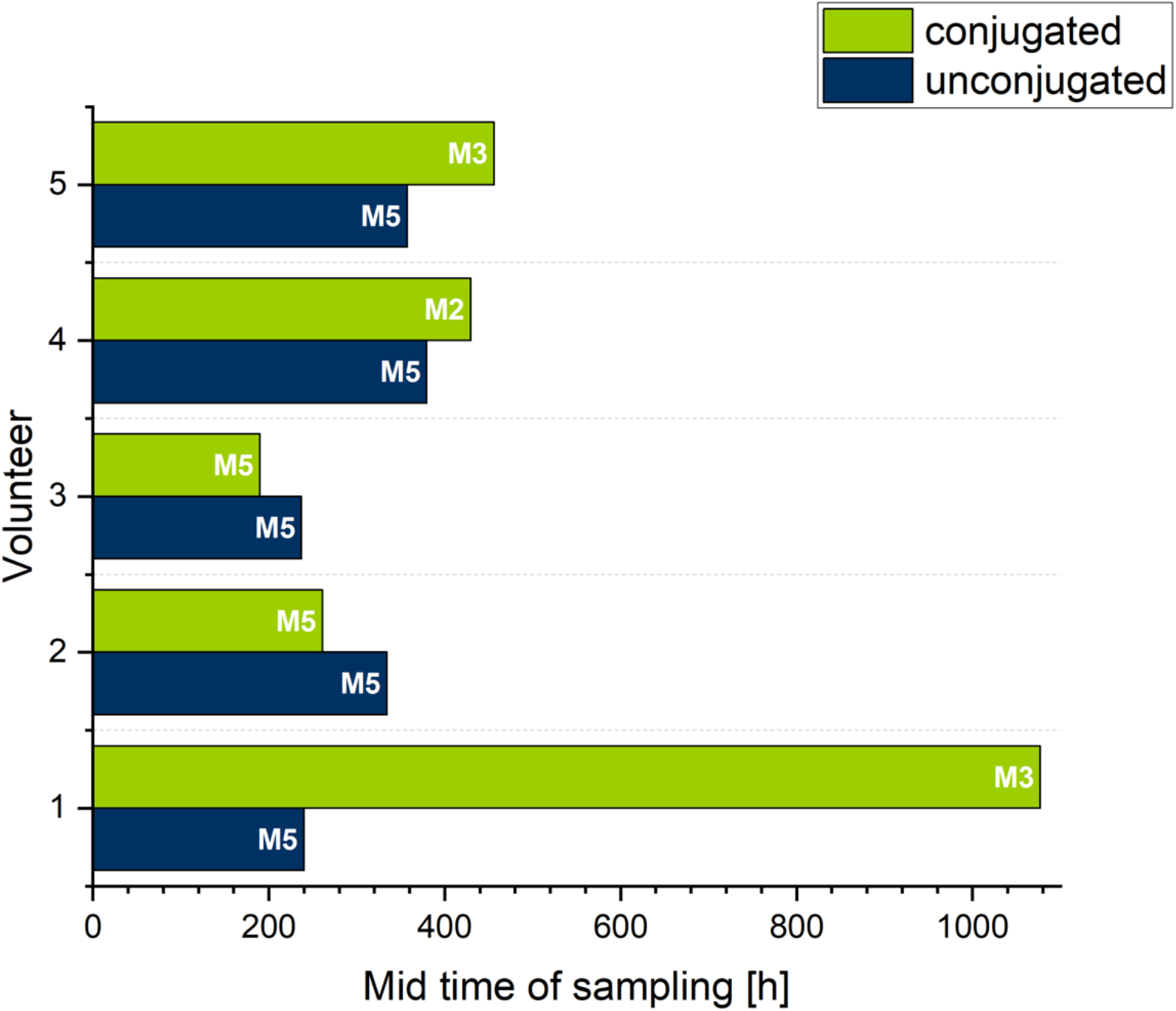
Comparison of the longest detectable metabolites in the free vs. the glucuronide fraction (all volunteers)

**Figure 9:**
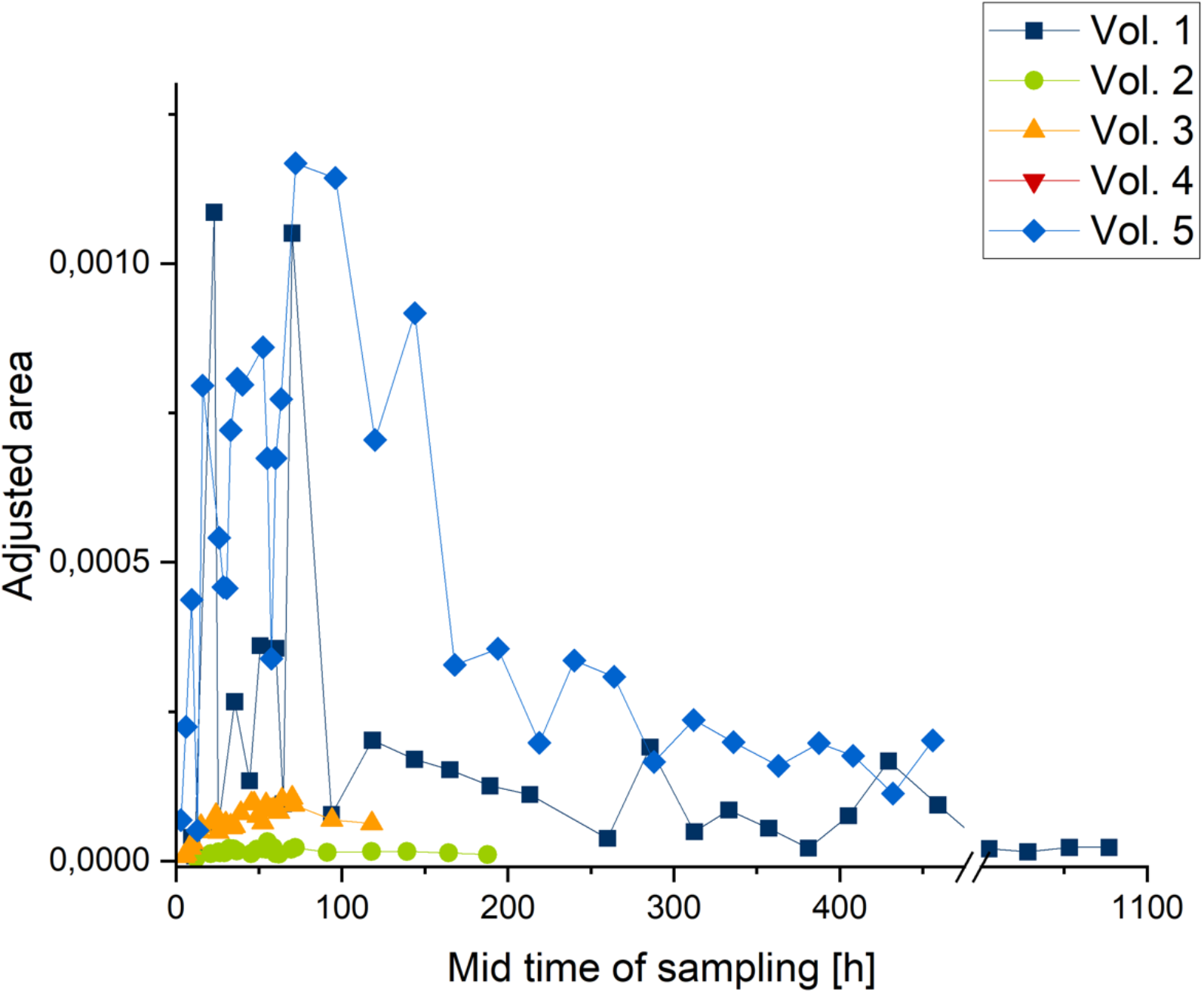
Elimination profile of metabolite M3 (detected in glucuronide fraction); peak area adjusted by internal standard and specific gravity

**Figure 10:**
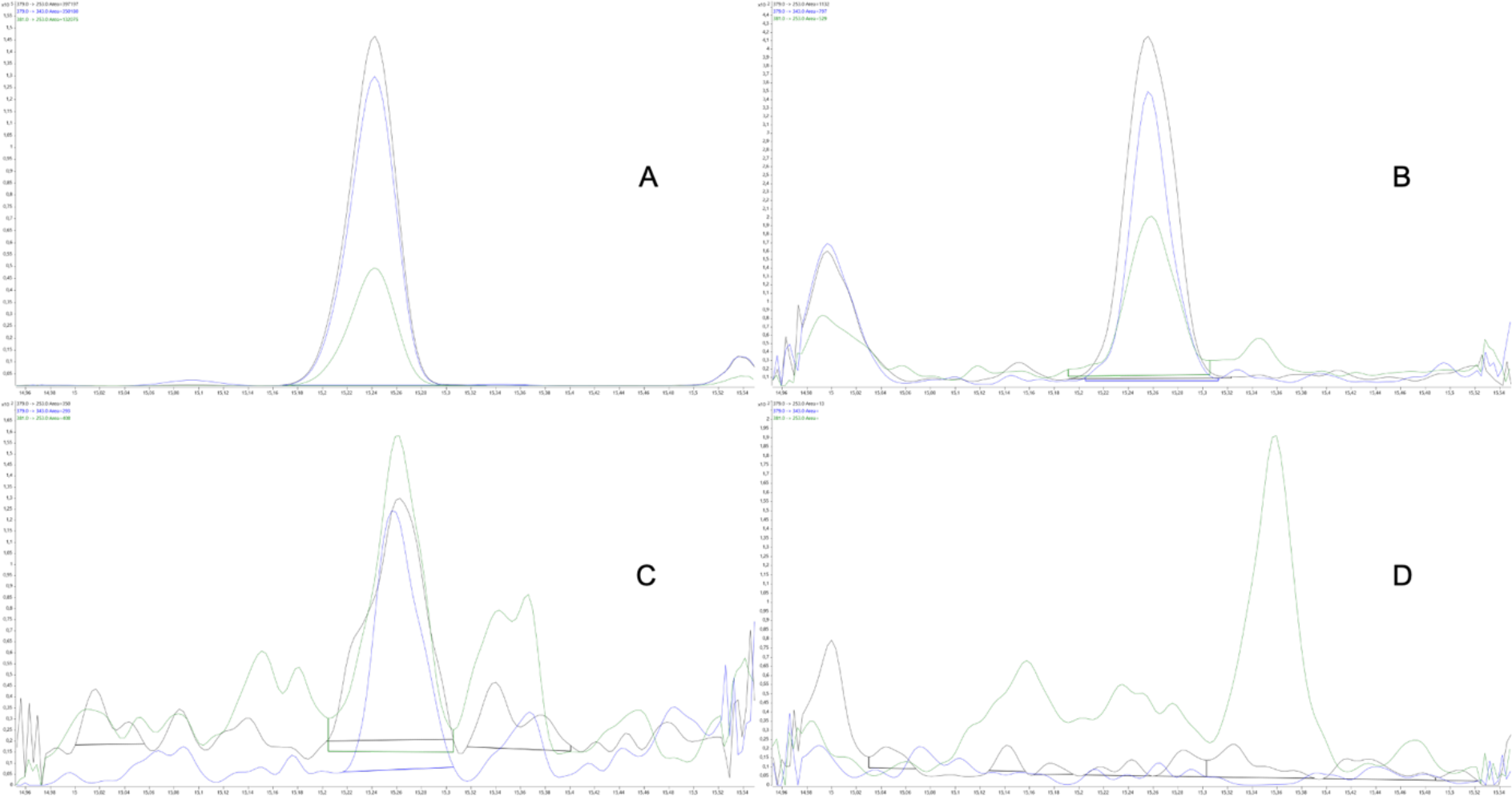
chromatograms (GC-QQQ-MS, 379→253, 379→343, 381→253) of volunteer 4: A standard M3 (1.8 ng/μL); B 24.1 h after intake; C 885.5 h after intake; D blank urine

The elimination profiles of the other metabolites (M1, M2, M4, epiM4, M5) are displayed in the supplement.

Excretion times after intake are displayed in Figure 11 and Figure 12.

**Figure 11:**
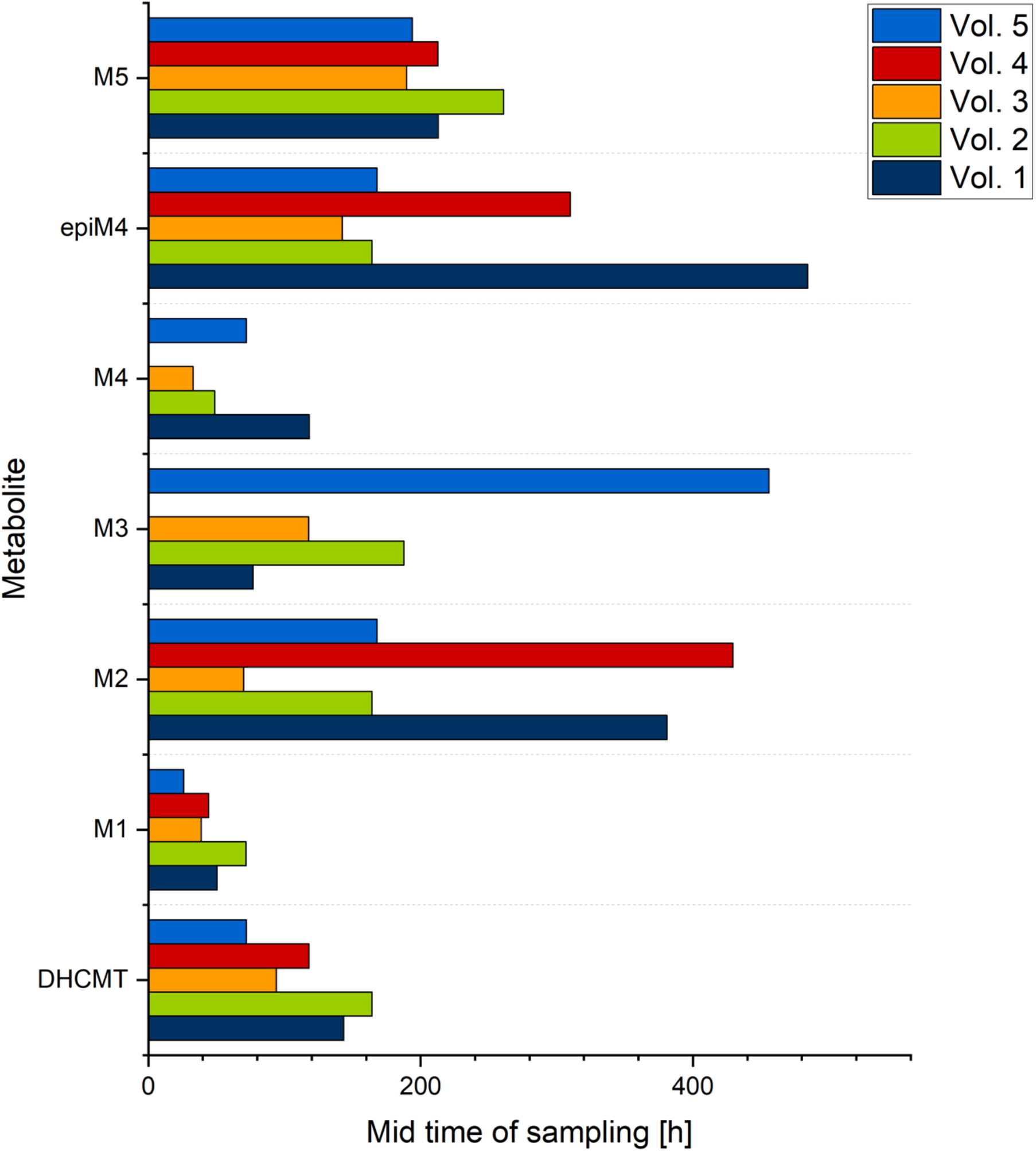
Last positive sample in the glucuronide fraction (time after intake in hours)

**Figure 12:**
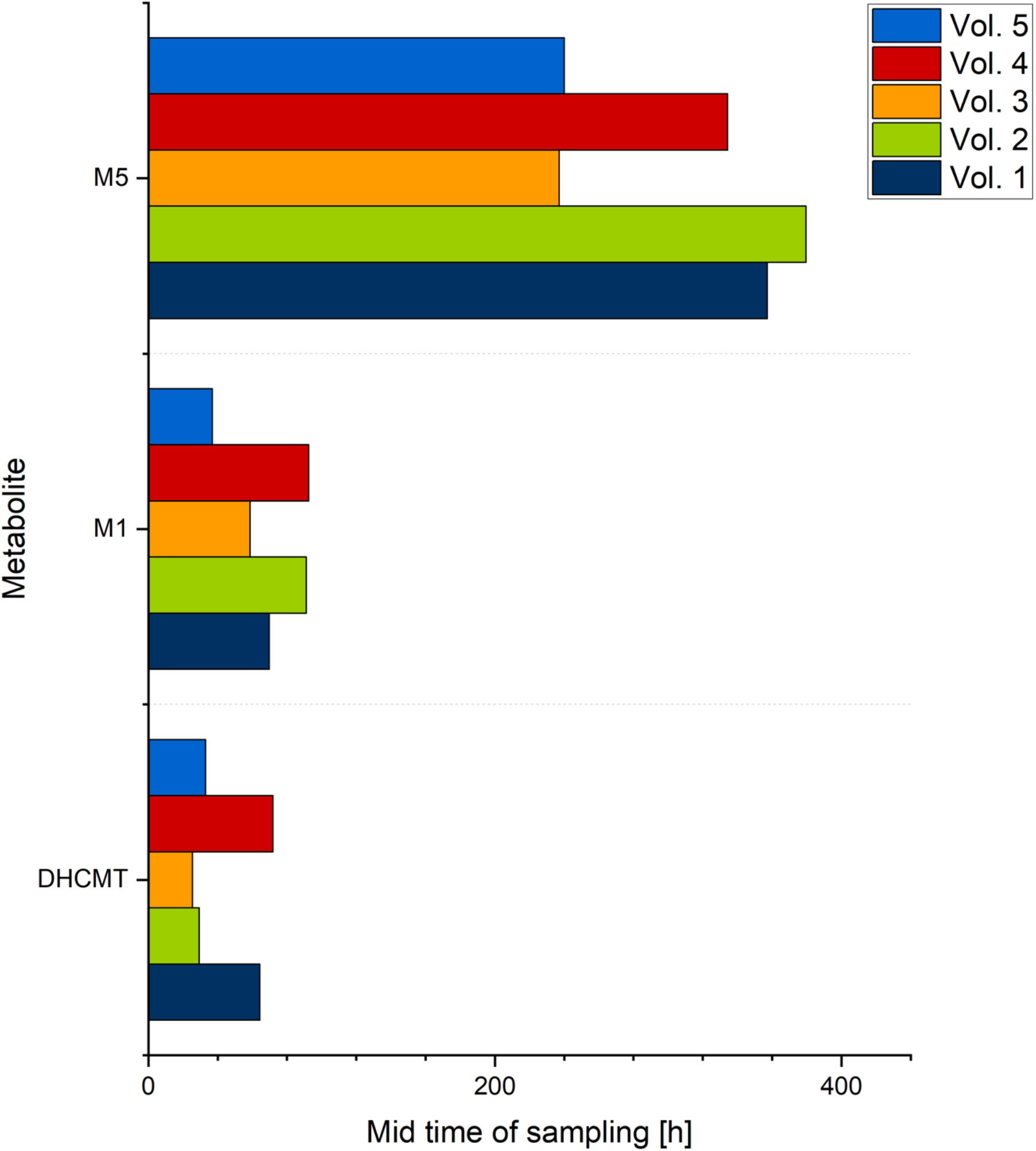
Last positive sample in the free fraction (time after intake in hours)

### Evaluation of Activities of Crucial Enzymes of Steroid Metabolism

For the calculation of the activity of the steroid 21-hydroxylase (CYP21A2), the areas of 3α5β-tetrahydrocortisone, 3α5β-tetrahydrocortisol, 3α5α-tetrahydrocortisol, pregnanetriol, and 17-hydroxypregnanolone in the blank samples were compared, as described by Krone *et al.* [40]. For evaluation of the activity of the 3-oxo-5α-steroid 4-dehydrogenase 2 (SRD_5_A2), the areas of etiocholanolone, androsterone, 3α5β-tetrahydrocorticosterone, 3α5α-tetrahydrocorticosterone, 3α5β-tetrahydrocortisol, and 3α5α-tetrahydrocortisol were used. The ratios are displayed in Table 4 and Table 5.

**Table 4:** Diagnostic ratios for diagnosis of inborn errors of steroidogenesis and steroid metabolism (steroid 21-hydroxylase) as used by Krone et al. [40]; 17HP: 17-OH-pregnanolone; THE: Tetrahydrocortisone; THF: Tetrahydrocortisol; 5α-THF: 5α-Tetrahydrocortisol; PT: Pregnanetriol

| Volunteer | Ratio 1<br>17HP / (THE + THF + 5αTHF) | Ratio 2<br>PT / (THE + THF + 5αTHF) |
| --- | --- | --- |
| 1 | 0.041 | 0.175 |
| 2 | 0.020 | 0.920 |
| 3 | 0.022 | 0.020 |
| 4 | 0.010 | 0.031 |
| 5 | 0.016 | 0.036 |

**Table 5:**
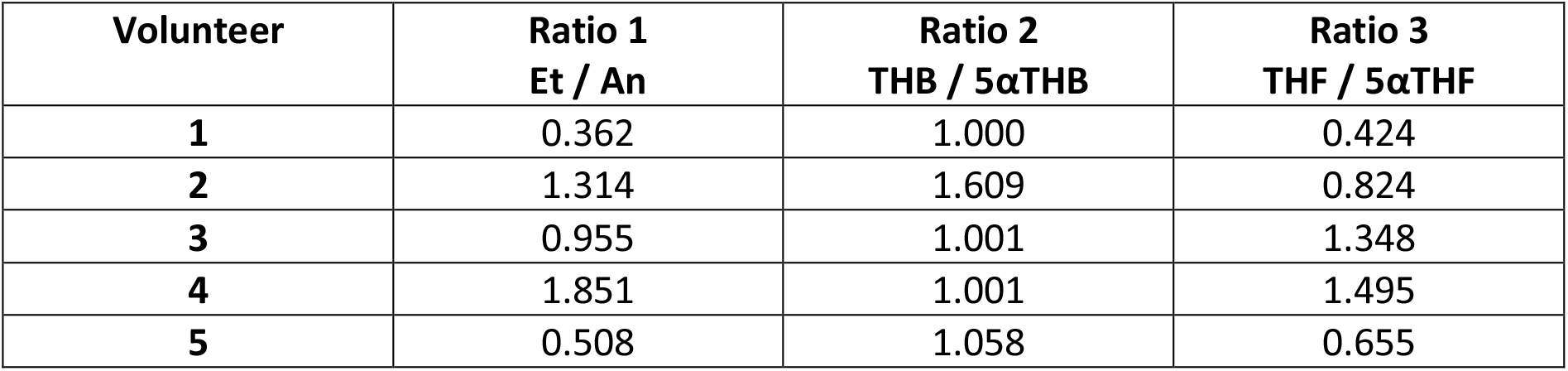
Diagnostic ratios for diagnosis of inborn errors of steroidogenesis and steroid metabolism (3-oxo-5α-steroid 4-dehydrogenase 2) as used by Krone et al. [40]; Et: Etiocholanolone; An: Androsterone; THB: Tetrahydrocorticosterone; 5α-THB: 5α-Tetrahydrocorticosterone; THF: Tetrahydrocortisol; 5α-THF: 5α-Tetrahydrocortisol

| Volunteer | Ratio 1<br>Et / An | Ratio 2<br>THB / 5αTHB | Ratio 3<br>THF / 5αTHF |
| --- | --- | --- | --- |
| 1 | 0.362 | 1.000 | 0.424 |
| 2 | 1.314 | 1.609 | 0.824 |
| 3 | 0.955 | 1.001 | 1.348 |
| 4 | 1.851 | 1.001 | 1.495 |
| 5 | 0.508 | 1.058 | 0.655 |

The displayed values for each volunteer are representing the ratios of the peak areas of the investigated compounds. For example, the ratio 1 of the 3-oxo-5α-steroid 4-dehydrogenase 2 is formed out of the areas of etiocholanolone and androsterone. When testosterone is metabolized, the first step is the reduction of the 4,5-double bond, giving the 5α- and the 5β-isomer. The following steps resulting in the above-mentioned compounds are catalyzed by the same enzymes for both ways (3α-HSD, 17β-HSD). Therefore, the ratios are giving an overview of the capacity of the 5α-reductase. A value above 1 means that the 5β-derivative is favored, and this is interpreted in a deficiency of the 5α-way.

## Discussion

All urine samples were analyzed by GC-QQQ-MS to meet the ultra-trace levels of metabolites. Due to the trace amounts of metabolites excreted in the urines, full direct identification was impossible, even after purification. Thus, GC-MS comparison with authentic reference material allowed for identification of DHCMT, 6-hydroxy-DHCMT and M3, resulting in level 1 confidence [41].

For confirmation of the tentative structures of M4/epiM4 no reference material was available. Thus, inhouse synthesis was required. Four diastereomeric androstane derivatives with a partially reduced A-ring (3ξ-hydroxy-4-ene) and a rearranged D-ring (17ξ-hydroxymethyl-17ξ-methyl-18-nor-13-ene) were obtained. As reported by Schänzer *et al.* [42] the reduction of 4-chloro-androst-4-ene-3,17-dione using K-selectride mainly results in the 3β-hydroxy isomer (ratio α/β ≈ 1/4). Due to the stereoselectivity of Wagner-Meerwein rearrangement the majority of C-17 substitution is hypothesized to yield 17α-hydroxymethyl-17β-methyl products.

After purification of the main product accurate mass spectrometry (GC-QTOF-MS) and NMR confirmed its structure as 4-chloro-17α-hydroxymethyl-17β-methyl-18-nor-androsta-4,13-dien-3β-ol.

Due to the extensive fragmentation caused by the high ionization energy of 70 eV no molecular ion was obtained. The dominant fragment [M-CH_2_-OTMS]^+^ (accurate mass *m/z* 377.2073, exact mass *m/z* 377.2067, Δ*m/z* = 1.59 ppm) was found. The loss of 103 Da is considered characteristic for per-TMS derivatives of 17-hydroxymethyl-17-methyl-18-nor-13-ene steroids. The fragment of *m/z* 287.1572 corresponds to another loss of TMSOH (exact mass *m/z* 287.1567, Δ*m/z* = 1.74 ppm). The fragments *m/z* 341.2304 and *m/z* 251.1800 are formed out of the above-mentioned fragments by loss of hydrochloric acid (exact mass *m/z* 341.2301, Δ*m/z* = 0.88 ppm; exact mass *m/z* 251.1800, Δ*m/z* = 0.00 ppm).

NMR data confirmed the structure proposal. Correlation analysis and multiplicity were used to assign 3-hydroxy as β-oriented. The related H-3α (3.92 ppm, ddm) showed a coupling with H-2β with ^3^J = 13.3 Hz, which may be explained by the pseudoaxial orientation. NOE of H-3α with H-2α (1.91 ppm, m) and H-1α (1.32 ppm, m) confirmed this assignment. The most shielded signal of C-17-CH_3_ detected at 0.90 ppm pinpoints towards β-orientation of the methyl group, concomitant with α-orientation of the 17-hydroxymethyl group. This assignment is also confirmed by correlation signals in NOESY. For confirmation of the substituent pattern at positions C-3 and C-4, predictions of the ^13^C chemical shifts for 4-chloro-17α-hydroxymethyl-17β-methyl-18-nor-androsta-4,13-dien-3β-ol and 3β-chloro-17α-hydroxymethyl-17β-methyl-18-nor-androsta-4,13-dien-4-ol were first calculated using the HOSE code in nmrshiftdb [43]: The observed and predicted shifts of C-3 are in good agreement for the expected OH group and C-4 for the expected chloro substituent. Utilization of high resolution ^13^C{^1^H} NMR yielded two signals for C-4 (130.47 ppm, 130.46 ppm). The second signal represents the carbon atom coupled to ^37^Cl isotope. No split signal was observed for C-3 (neither in 1D nor in HSQC), thus, confirming 3-hydroxy-4-chloro substitution [44, 45].

The side products were identified assigned based on the elution order (see Figure 13). As reported earlier [46], TMS derivatives of 3β-hydroxyandrost-4-enes elute later than their 3α-analogs. Similarly, 17α-hydroxymethyl-17β-methyl show shorter retention times than their 17β-hydroxymethyl-17α-methyl analogs [28]. Thus, the latest eluting isomer is assigned to 4-chloro-17β-hydroxymethyl-17α-methyl-18-nor-androsta-4,13-dien-3β-ol, the first eluting isomer is assigned to 4-chloro-17α-hydroxymethyl-17β-methyl-18-nor-androsta-4,13-dien-3α-ol. These assignments are also in line with the relative abundances of the isomers.

**Figure 13:**
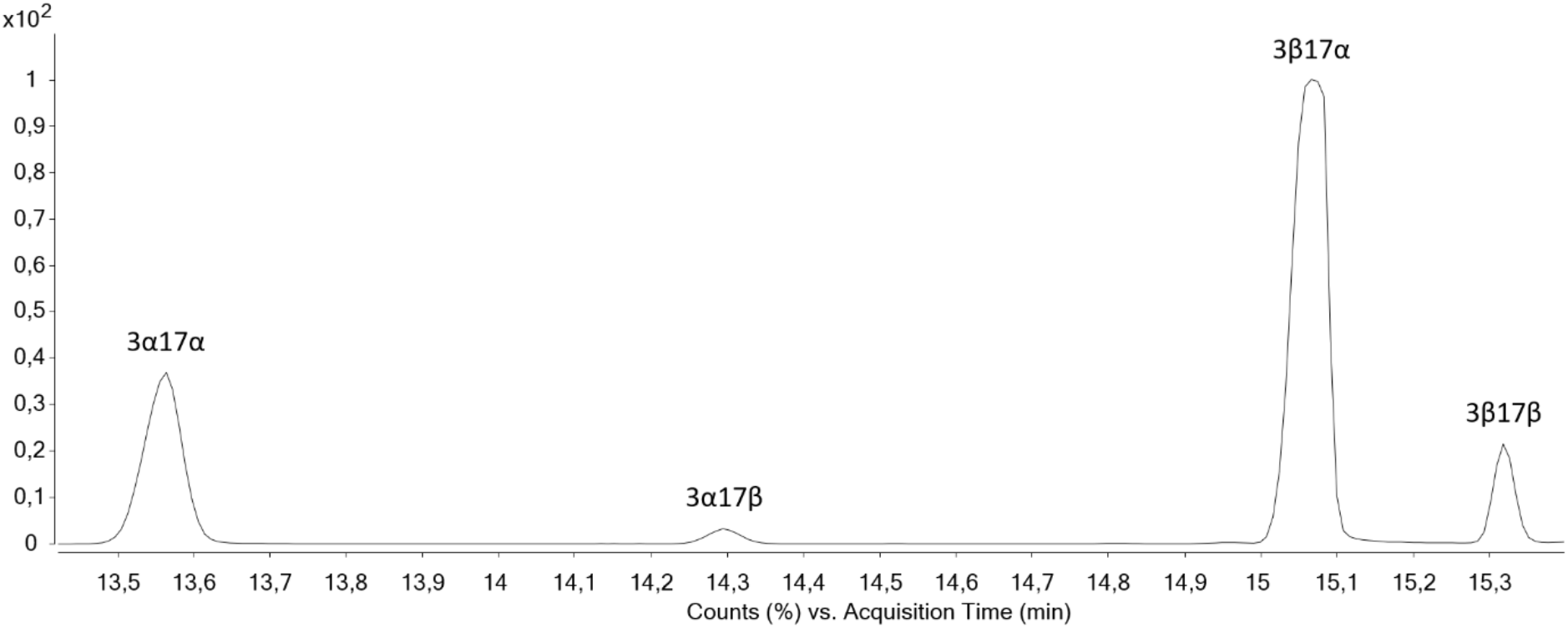
Elution order of the given isomers after synthesis (supposed stereochemistry)

GC-MS comparison of the synthesized references with the results obtained for the urine samples identified M4 and epiM4 as metabolites. Remarkably, that our structure assignments are challenging the hypotheses of Sobolevsky *et al.* in 2012 [23]. The DHCMT metabolite at the retention time of 15.1 min was assigned by Sobolevsky as 4-chloro-17β-hydroxymethyl-17α-methyl-18-nor-androstane-4,13-dien-3α-ol (3α17β). However, the isomer with this retention time is the main product from our synthesis (3β17α). For consistency in metabolite nomenclature, we call this metabolite epiM4, due to its 17α-hydroxymethyl structure. This data was confirmed by Kratena *et al.* [25]. Sobolevsky found an isomeric structure at a retention time of 14.3 min, which he called epiM4. He proposed a structure which is epimerized in position 17 (3α17α). The isomer assigned from our synthetic approach fitting this retention time has a configuration of 3α17β. Again, to be consistent in metabolite nomenclature we use “M4” for this 17β-hydroxymethyl metabolite. As reported [19, 28], the 17α-hydroxymethyl derivatives elute before their 17β-hydroxymethyl isomers. This is just the case if the rest of the molecule is identical. The observed structures in the study have a different A-ring conformation so this correlation does not apply. An overview of the isomer assignments is given in Table 6.

**Table 6:** Nomenclature of isomeric metabolites M4 and epiM4 in comparison to Sobolevsky’s nomenclature [23] including retention times

| Retention Time [min] | Nomenclature in this publication | Confirmed Structure (short form) | Nomenclature Sobolevsky 2012 | Hypothesized Structure (short form) |
| --- | --- | --- | --- | --- |
| 15.1 | M4 | 4-chloro-17β-hydroxymethyl-17α-methyl-18-nor- | epiM4 | 4-chloro-17β-hydroxymethyl-17α-methyl-18-nor- |
| | | androsta-4,13-dien-3α-ol<br>(3α17β) | | androstane-4,13-dien-3α-ol<br>(3α17β) |
| 14.3 | epiM4 | 4-chloro-17α-hydroxymethyl-17β-methyl-18-nor-androsta-4,13-dien-3β-ol<br>(3β17α) | M4 | 4-chloro-17α-hydroxymethyl-17β-methyl-18-nor-androsta-4,13-dien-3α-ol<br>(3α17α) |

However, based on well-known principles of phase II metabolism, it is likely that the 3-hydroxy group of the metabolites has an α-conformation as all examined substances are excreted as glucuronides. In contrast, steroids with a 3β-hydroxy group are mainly excreted as sulfates [15]. The subsequent investigation could focus on the sulfate fraction of the samples to verify this assumption. The glucuronidation of the hydroxy group at position 17 or 20 is unlikely because it is sterically hindered [15, 47]. Nevertheless, a glucuronidation of a 3β-hydroxy group or of the hydroxy groups in positions 17 and 20 might be possible.

The data reported here clearly show that it is not to monitor one single long-term metabolite, but to ideally screen for the entirety of all metabolites. This fact becomes apparent when considering the maximum detection time after intake of the substance: Two of the five volunteers excreted the metabolite M3 for the longest time (volunteers 1 and 5), in the samples of two other volunteers (volunteers 2 and 3), metabolite M5 was found for a longer time than any other. The last volunteer (volunteer 4) excreted metabolite M2 as longest detectable substance, while M3 could not be confirmed in any of these samples at all. Therefore, it is indispensable to track any use of DHCMT by screening for all these metabolites. Another crucial point is the combined examination of free and conjugated metabolites. Six of the eight targeted compounds are excreted over a longer period in the glucuronide fraction but for the mentioned metabolites M1 and M5 it is the other way round. The detection of M1 and M5 in the free fraction may be explained by fact that these metabolites are polar enough for being excreted directly after phase I metabolism without the need for conjugation to glucuronic acid. This is plausible due to the additional hydroxy group in position 6 (M1, M5) or the extra oxo group (M5). The calculated log P values are confirming this: log P (DHCMT) = 3.25; log P (M1) = 2.16; log P (M5) = 1.77. In a former investigation by Schänzer *et al.* [38] the metabolite M5 was found in the free fraction up to 9 days after intake of 40 mg of DHCMT. This goes in line with the findings from our study.

The common practice in anti-doping analysis is the cleavage of the glucuronides and analysis of the combined free and glucuronide fraction. This is suggested to be a reasonable way to target all metabolites in one analytical run. Furthermore, especially relevant for DHCMT, M1, and M5, this procedure increases the concentration in the final extract, which may further prolong their detectability.

Sobolevsky *et al.* [23] estimated the excretion time of the metabolite M3 to 40-50 days. In volunteer 1, we indeed found the proposed long-time marker up to 45 days. However, due to the samples that had different signals than the blank urine but were not clearly identified as positive (e.g., lack of single transitions, different ratios of transitions to each other), there is a chance that this window can be further extended by starting from a greater volume of urine (> 6 mL).

For many drugs or xenobiotic substances in general, the body’s distribution is displayed by the Bateman function (Figure 14**Fehler! Verweisquelle konnte nicht gefunden werden.**). We assumed idealized one-compartment body model with first-order absorption and elimination as a basis [48]. A first-order kinetics is characterized by the fact that an even proportion of the substance is absorbed or eliminated per time unit [49, 50]. Based on the data from our analyses, especially the two or more cases of increasing concentrations of metabolites after a period of decreasing concentrations (cf. excretion of parent compound of volunteers 1 and 5), it is highly unlikely that the kinetics of DHCMT, i.e., absorption, distribution, metabolism, excretion, follow a first-order kinetics. Therefore, it cannot be easily described by the Bateman function. Another fact that hampers a smooth excretion curve is the potential effect of enterohepatic cycling. This is well described for estrogens but not investigated for androgens [51]. Another compounding factor could be the different time interval of each days’ urine collection: The volunteers did not orient themselves toward the collection process, so on every day there is a variable period between the last urine in the evening that was discarded and the saved one in the morning: this and the minimal concentration of the analytes exacerbate the identification of a defined excretion order.

**Figure 14:**
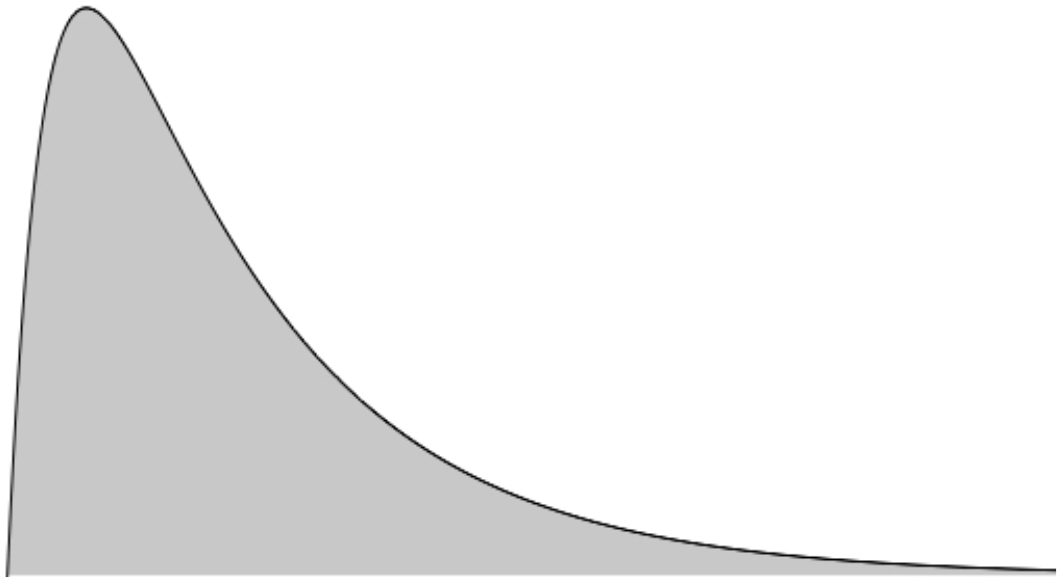
Bateman function; general form: 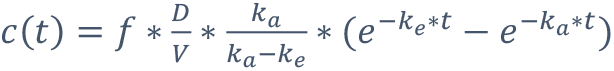

Based on our findings and the common metabolism of anabolic-androgenic steroids, we propose the following mechanisms in the formation of the metabolites investigated in this study.

M1: CYP3A4 hydroxylates testosterone in position 6 [52, 53]. Due to the structural similarity, Rendic *et al.* demonstrated that 6β-hydroxylation also occurred *in vitro* after incubation of DHCMT with recombinant CYP3A4 while CYP2C9 or CYP2B6 did not catalyze this reaction in DHCMT. The stereoselectivity of the catalyzed reaction is explained by the electronic effects of 3-keto-4-ene steroids through the stabilization of the substrate-radical intermediate formed during the reaction [54].

M2-M4: The formation of the altered D-ring structure (17β-hydroxymethyl,17α-methyl-18-norandrost-13-ene) starts with the sulfonation of the 17β-hydroxy group, followed by elimination of sulfuric acid, and a Wagner-Meerwein rearrangement [55]. The hydroxylation of the 17β-methyl group follows catalyzed by CYP21A2 [19]. This results in the intermediate generation of 4-chloro-17β-hydroxymethyl-17α-methyl-18-norandrost-1,4,13-trien-3-one [27]. For generation of the A-ring reduced metabolites, these steps are required to happen first because the hydroxylation of the 17β-methyl group only takes place in the presence of a 3-oxo function. The hydroxylation by CYP21A2 will not take place in the case of 3-hydroxy steroids [29]. An alternative pathway towards this intermediate may be the hydroxylation of C-18 by CYP11B2, followed by a Wagner-Meerwein rearrangement, which was demonstrated for metandienone [28]. Investigations on structure requirements for these reactions should be carried out in the future.

M2: As described by Schänzer, the initial and rate-limiting step in the reduction of the A-ring is the reduction of the 4,5-double bond, directly followed by the reduction of the 3-oxo group. The first step is likely catalyzed by the 5β-reductase as the chlorine atom in position 4 as well as the double-bond in position 1 hamper 5α-reductase (no 5α-metabolite reported for 3-oxo-1,4-diene steroids except M3). The second step is most likely catalyzed by the 3α-hydroxysteroid dehydrogenase. No 3β-formation in a 5β-steroid was ever reported in humans and 3α-steroids are reported to be mainly excreted as glucuronides, which is also the case for M2 [15].

M3: The structure assignment by Forsdahl *et al.* is 3α-hydroxy-4α-chloro-5α. It is remarkable that the A/B-ring connection is *trans* (5α), while the other reduced metabolites of DHCMT show likely 5β-orientation. Recent data of our group elucidating the structure of new metabolites of metandienone and methyltestosterone including a proposed order of reductions in the A-ring [56] suggest that the double bond in position 1 is reduced before the one in position 4 (cf. supposed formation of M2). Subsequently, the reduction of the 4,5-double bond and of the 3-oxo group take place. Nevertheless, the reduction of the 1,2-double bond is described for metandienone in the presence of a 3-hydroxy group [57]. Massé *et al.* considered a different order in A-ring reduction of metandienone to be possible (first reduction of the 1,2-double bond, then of the 4,5-double bond / 3-oxo group). Unfortunately, no proof of this assumption is published until now [58].

M4: For M4, the next steps are the reductions of the 3-oxo group by 3^α^-hydroxysteroid dehydrogenase (3α-HSD) and of the 1,2-double bond.

M5: The hydroxylation in position 6 is described above for M1. The introduction of the oxo-function in position 16 can be a product of 16-hydroxylation by CYP3A4, followed by oxidation. 16-Hydroxylation is well-known in the metabolism of estrogens [59] but also reported for testosterone [53]. The enzyme responsible for the following step, the oxidation of the 16-hydroxy group, remains unclear for now, whereas the steps afterward (reduction 4,5-double bond, reduction 3-oxo group) are catalyzed as described above.

The formation of epiM3 and epiM4 is not elucidated yet. For the case of epiM3 no sample was clearly marked positive.

In case of the metabolites M2, M3, M4, and M5, the formation is hypothesized. Further experiments for confirmation are desired in the future. An overview of the postulated metabolism of DHCMT is displayed Figure 15.

**Figure 15:**
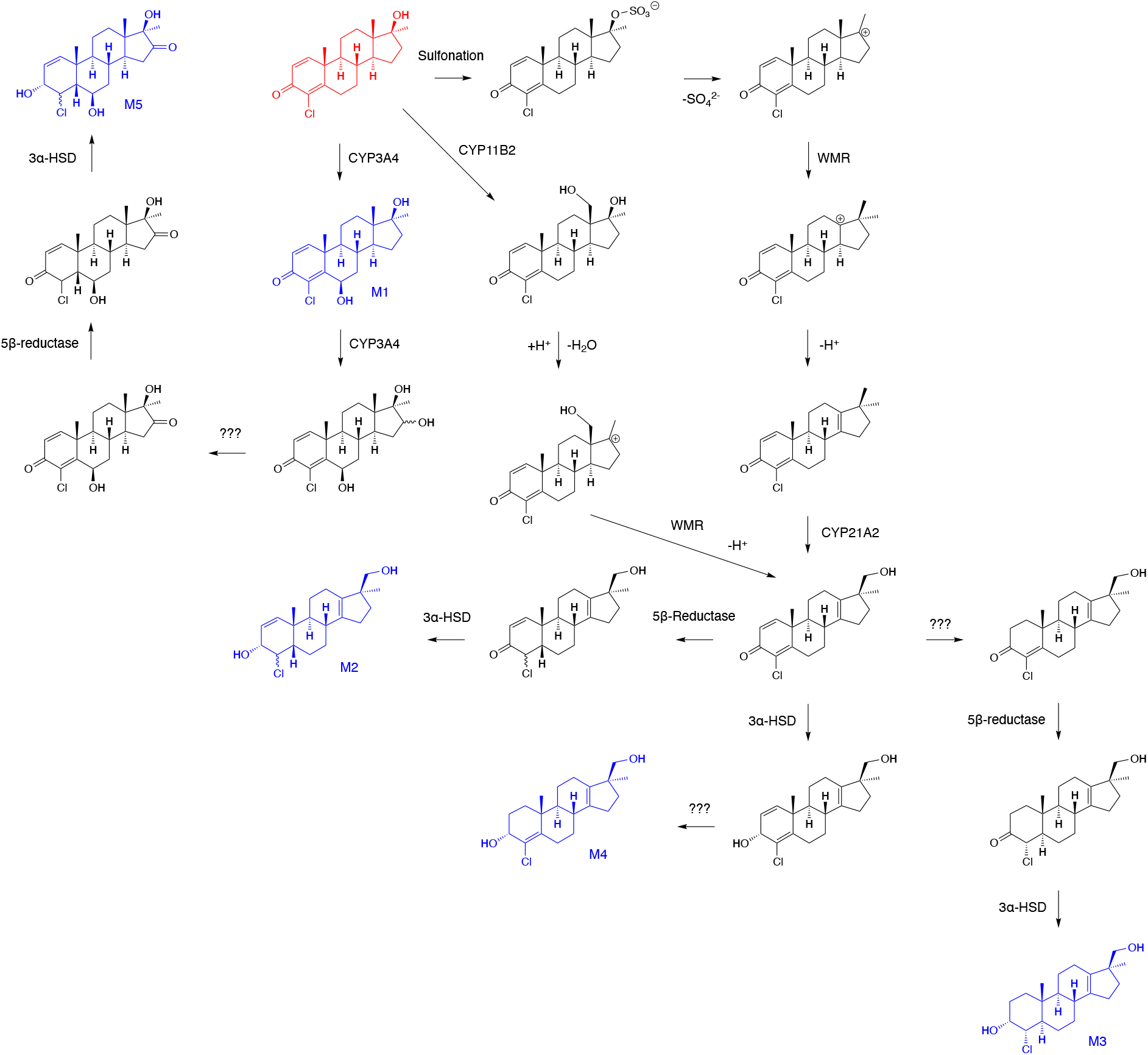
Postulated metabolism of DHCMT including the formation of metabolites M1-M5; question marks indicate enzymes that have not been elucidated yet; red = parent compound; blue = targeted metabolites

The differences in the excretion of the long-time markers M3 and M4 between the younger volunteers (vol. 1, 2, 3, and 5) and the older one (vol. 4) may be explained by the proposed metabolic pathways leading to these metabolites. As M2, M3, and M4 supposably share their D-ring structure, it is assumed that the difference in pathways of formation happens later. From the data of the extended steroid profiles all volunteers show a sufficient activity of CYP21A2. It remains unclear why volunteer 4 excretes epiM4 and M2 but not M3/M4. Therefore, a deficiency in the enzyme that reduces the 1,2-double bond is not plausible. The metabolite epiM3 was not clearly identified in any sample. This could have been caused by the fact that the other ways of metabolite formation are highly favored in comparison to epiM3. Other possibilities are a different location of the involved enzymes or a kinetic problem. Due to the fact that the metabolism of 17-epi metabolites of DHCMT is not elucidated yet, these occurrences cannot be explained for now.

The activity of 5α-reductase and its impact on the formation of different metabolites is hard to evaluate, because the finally excreted amount of metabolite is caused by the complex metabolism. Additionally, the structures of the metabolites M2, epiM3 and M5 are not confirmed yet due to the lack of reference material. Nevertheless, the values can provide some interesting information regarding the structure of the metabolites. Volunteer 4 excretes the metabolite M2 longer than any other volunteer. At the same time, he has the lowest 5α-reductase activity. This could provide an indication of the structure of this metabolite and favor the 5β-structure. If the volunteers 2 and 3 had higher 5α-reductase activity, perhaps they would produce more M3 and thus excrete it longer. Because of the almost independent excretion of metabolite M5 of the 5α-reductase activity, this indicates that the structure assignment of Sobolevsky was right (5β).

Finally, the various differences in metabolite formation in combination with the excreted amount of parent compound underline the fact that anabolic-androgenic steroids are extensively metabolized and that the location and point in time of metabolism are mostly unknown.

## Conclusion

The controlled administration study confirmed that the analytes proposed by Sobolevsky *et al.* can be used to trace back the intake of DHCMT. Our findings support Sobolevsky’s hypothesis of a long-term metabolite with a reduced A-ring and an 18-nor-13-ene-17α-methyl-17β-hydroxymethyl structure. These findings support the current practice of routine doping analysis by targeting these metabolites.

In routine anti-doping control, an aliquot of 2 mL urine is prepared prior to analysis combining the unconjugated and glucuronidated metabolites. In this study 6 mL of urine have been used to enlarge the detection time of the metabolites. Separate analysis of the phase-II metabolite fractions gave further insights into the metabolism. The results show that some metabolites can be identified clearly until a certain point of time but are likely excreted even longer. Still, higher amounts of urine, such as 20 mL, might be used to possibly further extend the detection window of this metabolite. Due to the difficulties in handling such a large amount of sample volume, this procedure may not be suitable for routine analysis but could be interesting in confirmatory testing or in future studies of metabolism.

In this study, all volunteers belong to the Caucasian ethnicity. Nevertheless, the excretion profiles of the different volunteers are considerably different. Therefore, it is indispensable to monitor all metabolites irrespective of the expectation of the longest detectable metabolite. To verify the proposed metabolite structures, it is mandatory to synthesize reference material to achieve the highest confidence level [41]. A level 1 confirmation of the structures of M2 and M5 would be highly desirable.

A potential next experiment will be the inclusion of volunteers from other ethnic groups to investigate potential different metabolism pathways of non-Caucasian test persons. The determination of characteristics (polymorphism, dysfunctionality, et cetera) of the future volunteers’ enzymes like UDP-glucuronosyltransferases or cytochromes P450 would be helpful. Additionally, blood samples and tissue could be collected during the next study to calculate the volume of distribution and to uncover potential drug and metabolite depots. This may help in understanding the distribution of the substances.

To elucidate the remaining ‘blind spots’ in DHCMT metabolism, further experiments with potential intermediate metabolites and purified enzymes are mandatory. Parallel investigations on the substrate selectivity using molecular modeling techniques may explain and substantiate the experimental data.

## Acknowledgments

We would like to thank the World-Anti Doping Agency (WADA) for their project support (WADA 17C02MP).

