## Supplemental Material for "Controlled Administration of Dehydrochloromethyltestosterone in Humans: Urinary Excretion and Long-Term Detection of Metabolites for Anti-Doping Purpose"

### Synthesis of 4-Chloro-17α-hydroxymethyl-17β-methyl-18-nor-androsta-4,13-dien-3β-ol

A 0.05 M solution of 4-chloro-androst-4-ene-3,17-dione in dry $Et_2O$ under argon atmosphere has been treated with 1.1 eq of K-selectride (1 M) by dropwise addition to the stirring solution. After 1.5 h the reaction mixture was diluted with water and extracted three times with $Et_2O$. The combined organic phases were washed with brine and dried with $Na_2SO_4$. Evaporation at reduced pressure gave the crude compound as a yellow solid.

The reduced substance was dissolved in dry DMF (0.1 M) then 2.5 eq of imidazole and 1.5 eq of TBSCl have been added at room temperature. After 48 h of stirring the solvent was evaporated. The crude solid has been solubilized in EtOAc/$H_2O$ and transferred in a separatory funnel. The obtained organic phases have been washed with brine and dried with $Na_2SO_4$. After evaporation the product was obtained as a yellow oil.

A solution of Nysted reagent (20 % wt., 5 eq) diluted in half volume of dry THF was cooled to 0 °C and subsequently treated with 2.2 eq of $TiCl_4$. After stirring for 15 minutes at 0 °C the solution was allowed to warm to room temperature. Next, a solution of starting material dissolved in dry THF was added dropwise to the reaction flask to give a violet mixture. After overnight reaction at room temperature the reaction was quenched with $HCl_{aq}$ (1 M) at 0 °C. Following dilution with water and the mixture was extracted three times with $Et_2O$. The combined organic phases were washed with brine and dried over $Na_2SO_4$. Evaporation at reduced pressure gave the crude compound as a yellow solid.

To a 0.1 M solution of the olefin in DCM, 3 eq of $KHCO_3$ and 1.2 eq of m-CPBA have been added at room temperature. After 3 hours, the solution was diluted with water and extracted with DCM in a separatory funnel. The combined organic phases were washed with brine and dried with $Na_2SO_4$, then evaporated to give the crude compound as a yellow solid.

To a solution 0.04 M of DCM under argon atmosphere at -78 °C, 2.2 eq of 2,6-lutidine and 2 eq of TMSOTf have been added dropwise. After five minutes the compound has been solubilized in DCM and subsequently added dropwise to the reagent’s solution at -78 °C. After one hour the reaction mixture has been quenched by the addition of $HCl_{aq}$ 2 M (10 eq) and let stir for 30 minutes at room temperature. Then, the solution was diluted with water and extracted with DCM in a separatory funnel. The combined organic phases were washed with brine and dried with $Na_2SO_4$, then evaporated to give yellow mixture.

To a solution 0.04 M of epoxide in MeOH, 5 eq of aqueous HCl have been added dropwise. After 1 hour the Wagner-Meerwein rearrangement and deprotection was already accomplished. The mixture was let react overnight at room temperature. Then it has been quenched by addition of water and diluted in TBME and extracted in a separatory funnel. The combined organic phases were washed with brine and dried with $Na_2SO_4$ then evaporated to give the crude final product as a yellow oil.

Supplement S2
Synthesis of epiM4 isomers

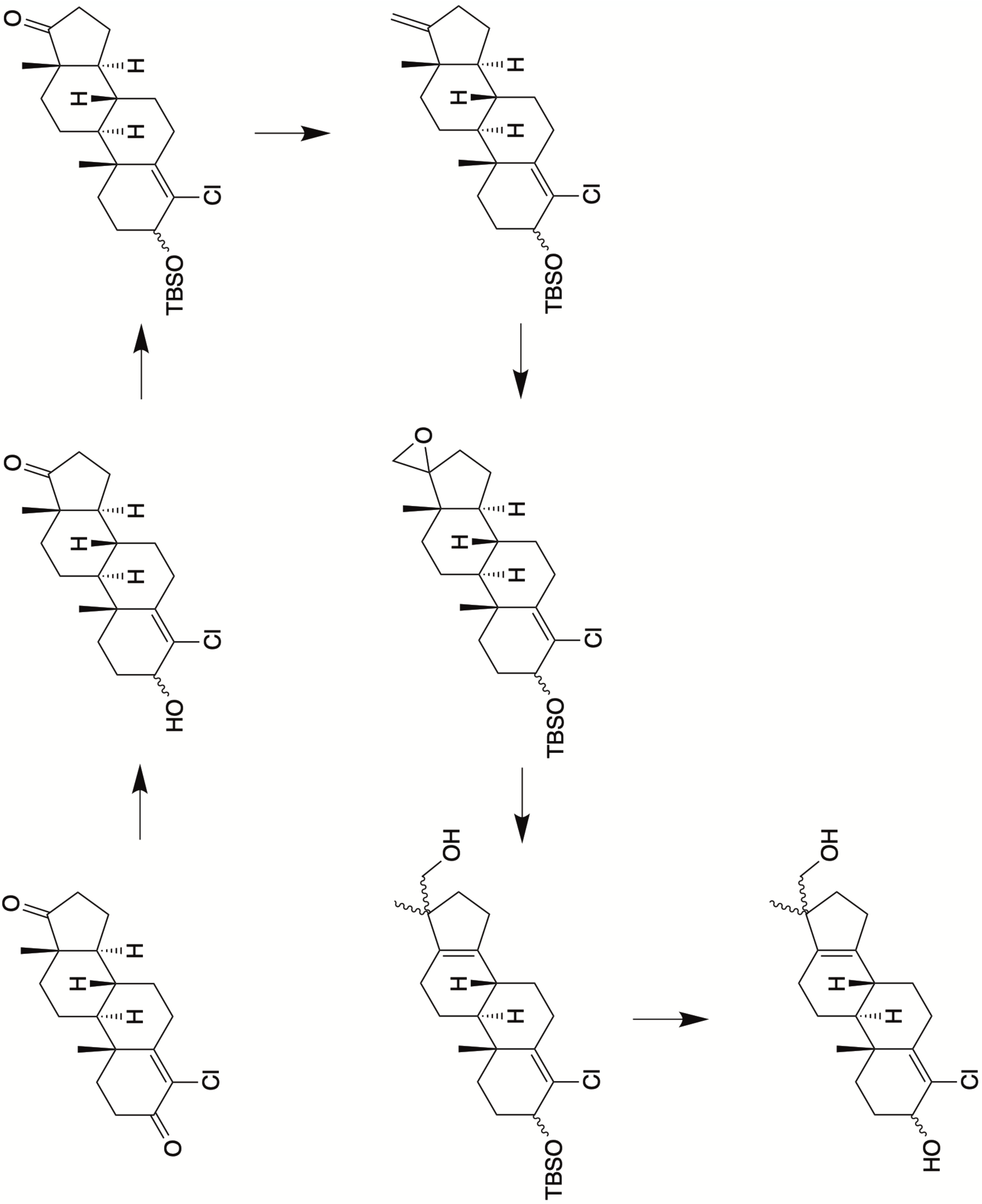

Supplement S3
Excretion profile M1 (glucuronide)

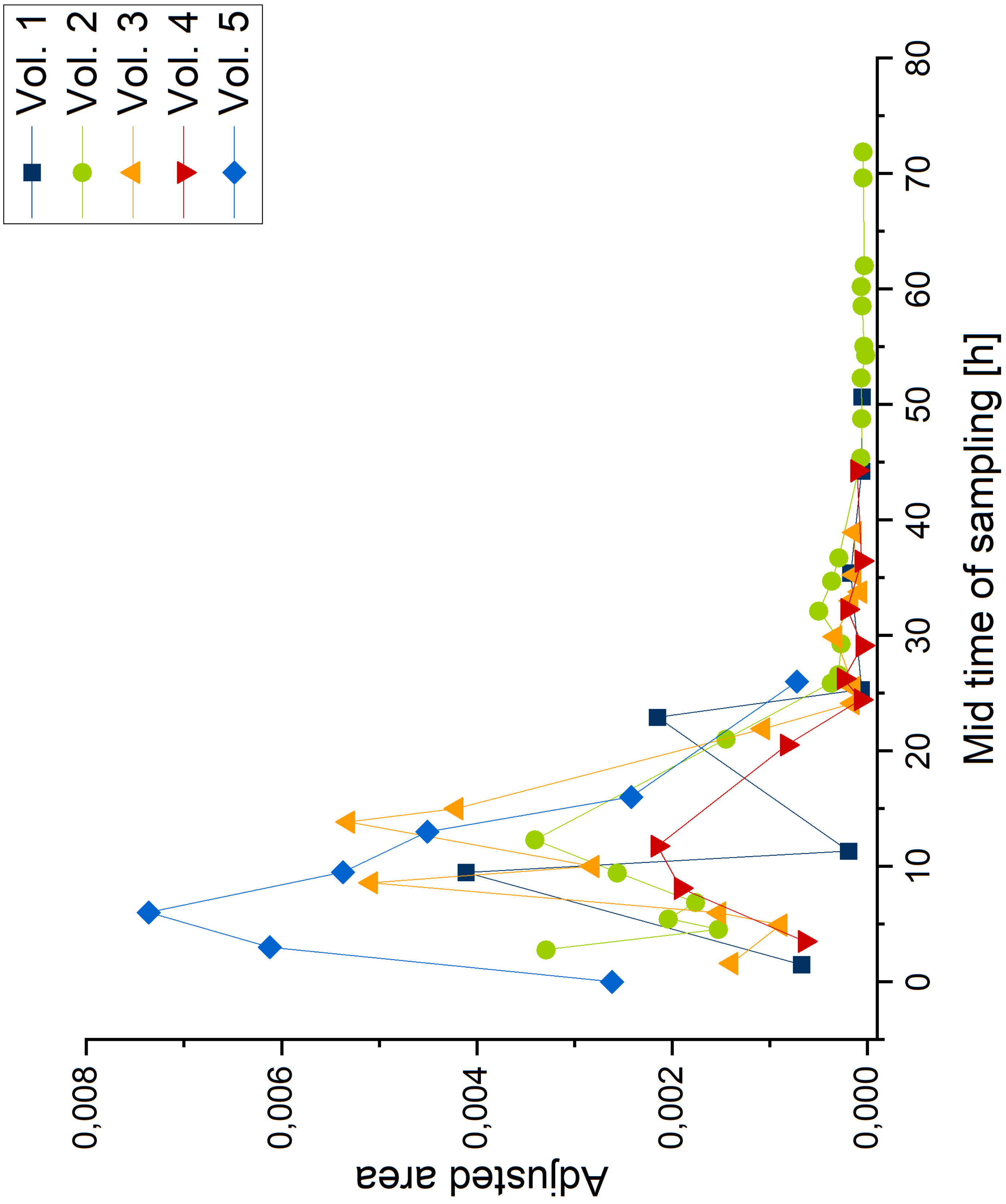

Supplement S4

Excretion profile M2 (glucuronide)

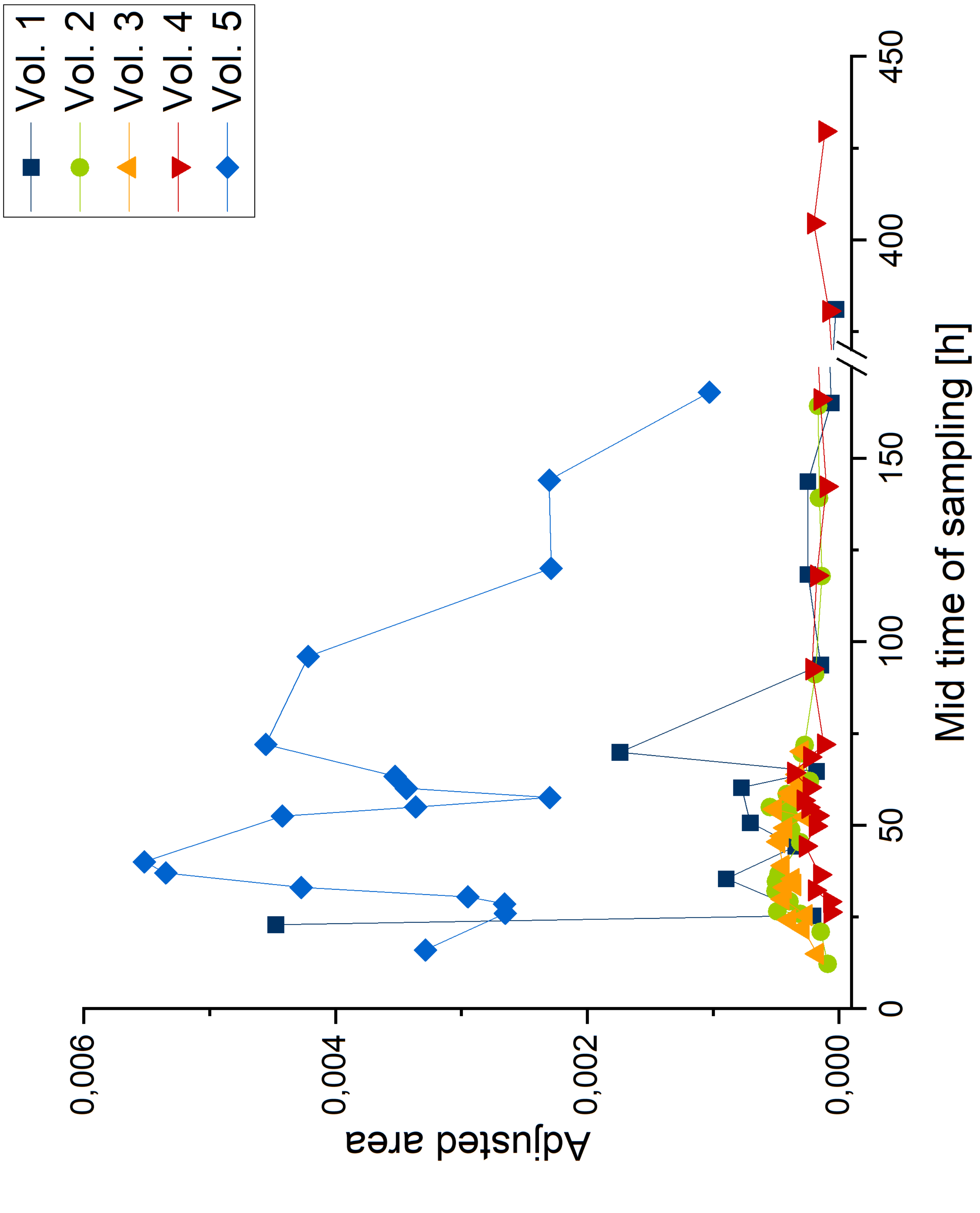

Supplement S5
Excretion profile M4 (glucuronide)

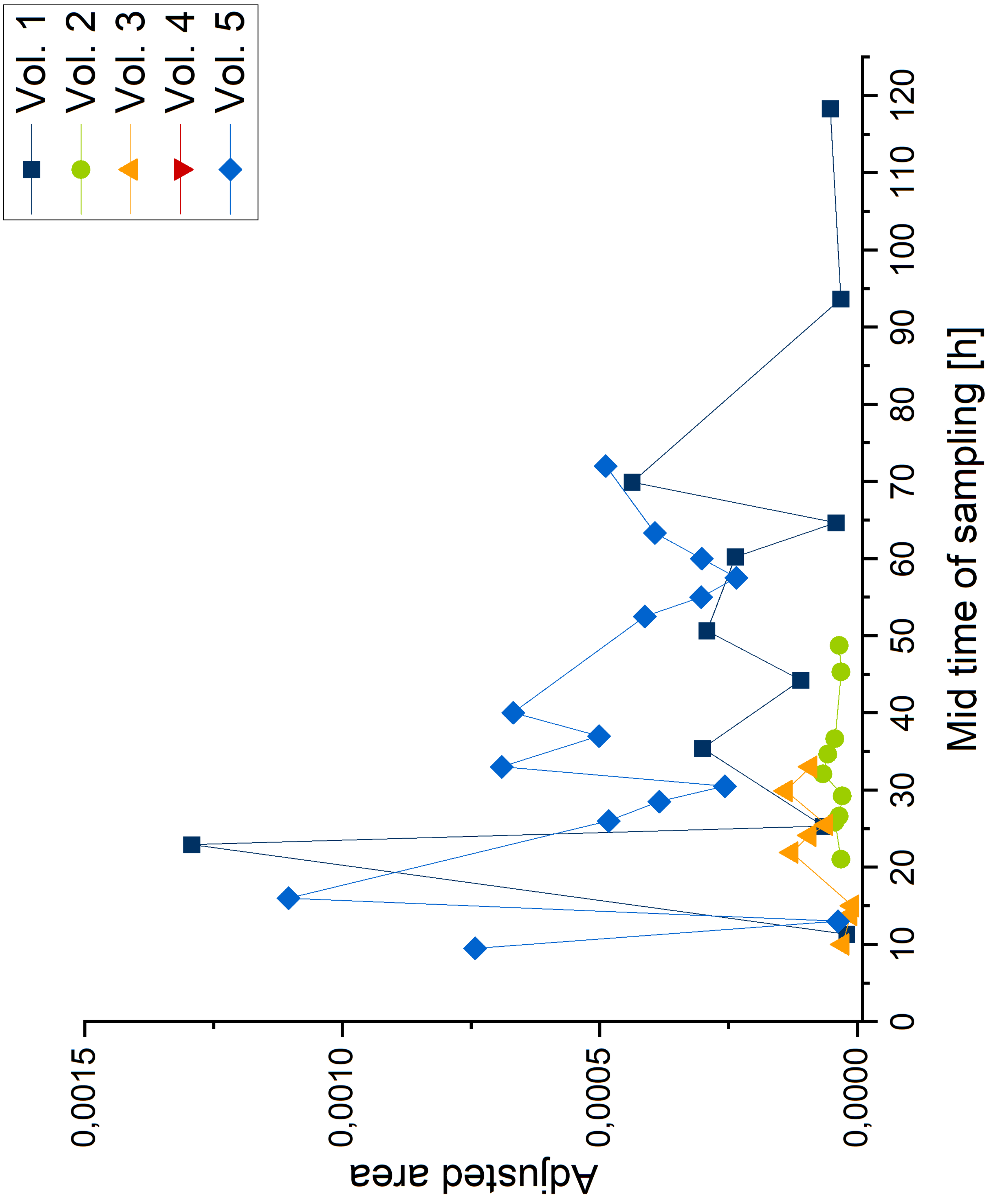

Supplement S6
Excretion profile epiM4 (glucuronide

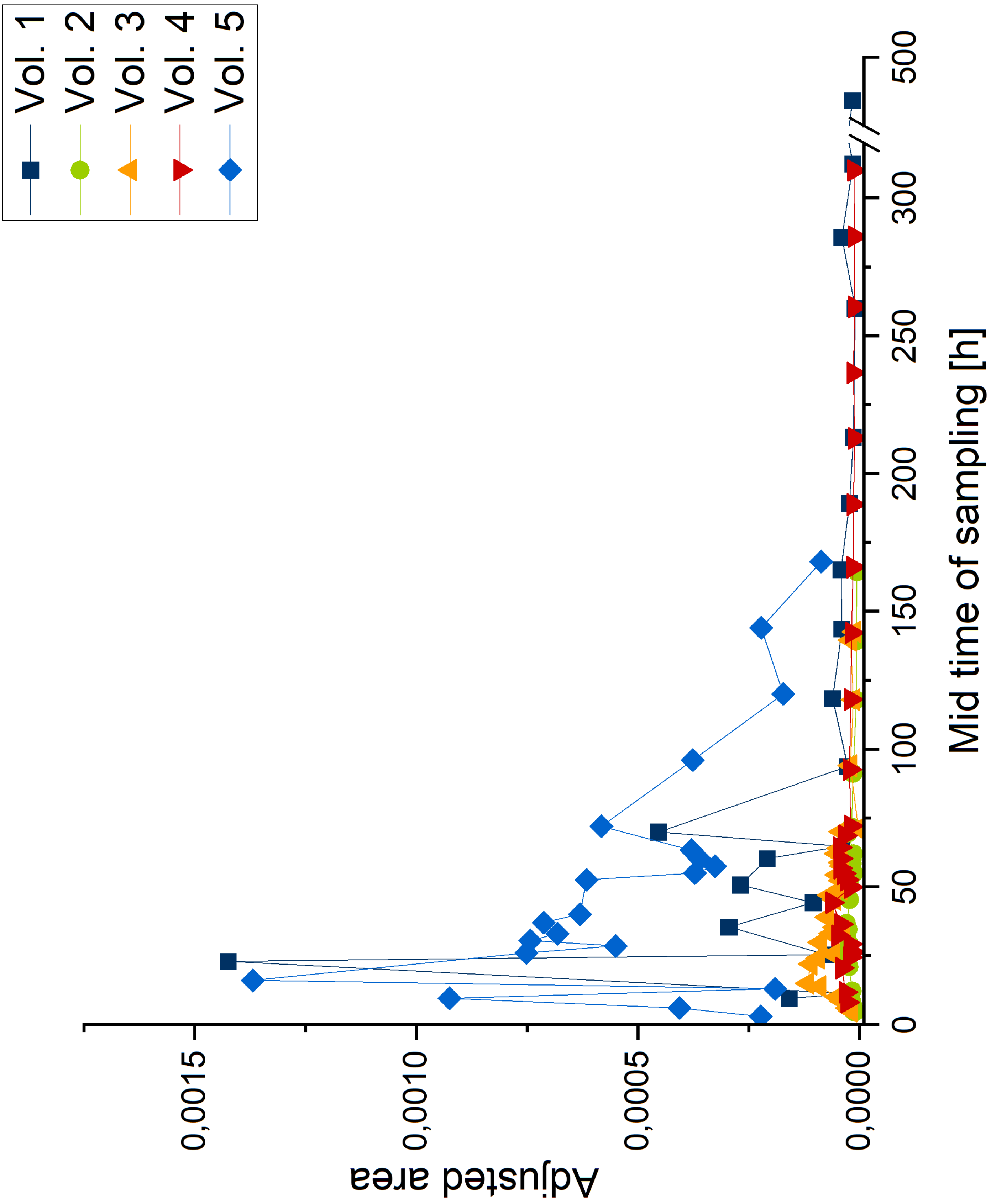

Supplement S7
Excretion profile M5 (glucuronide)

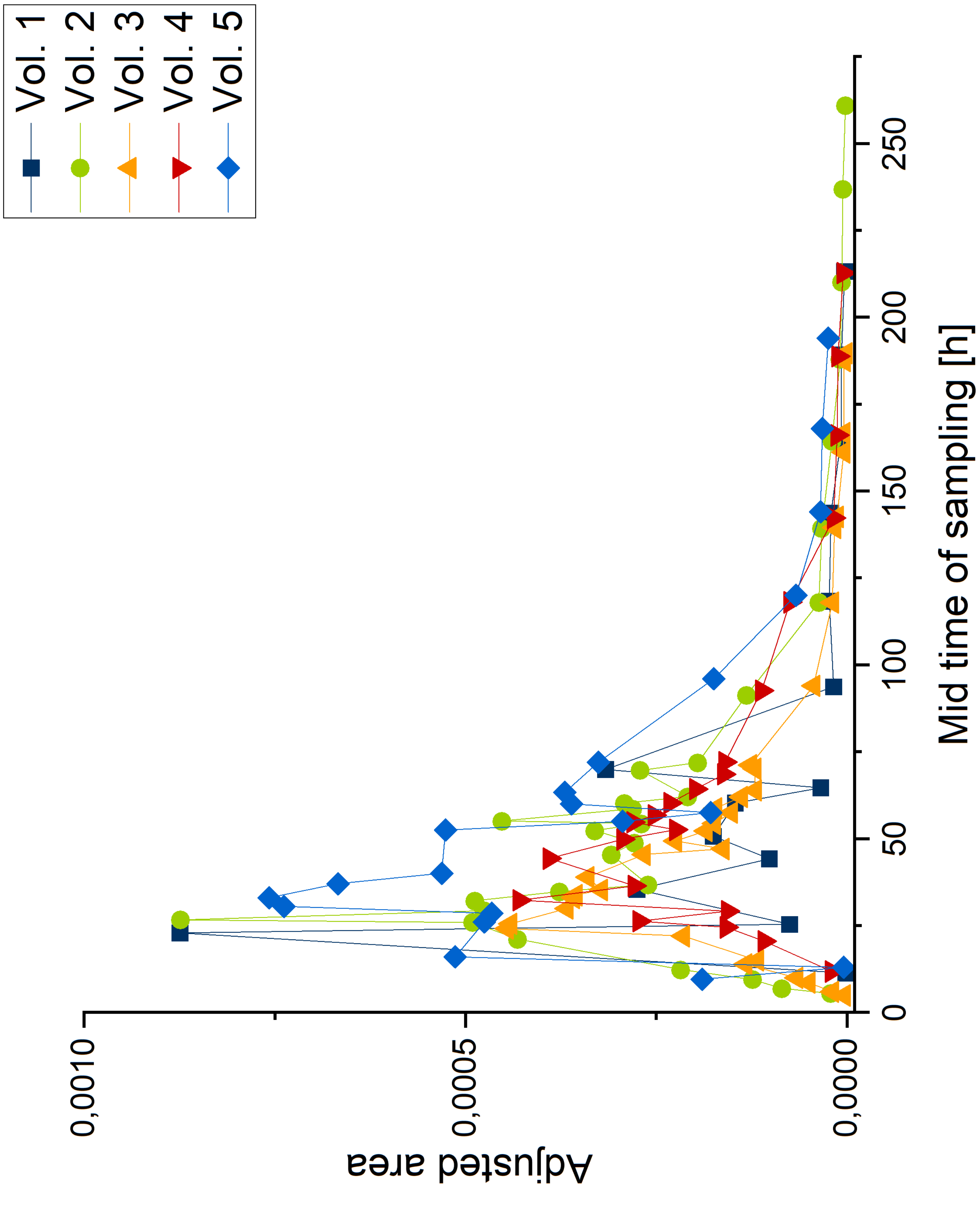
